# A Nanomechanical Fingerprint of Centromeric Methylation Co-Localization in TNBC Chromosomal Instability

**DOI:** 10.64898/2026.10.01.756136

**Authors:** Tanya Agrawal, Shoubhik Chakraborty, Sejal Khanna, Sarthak Sahoo, Mohit Kumar Jolly, Sneha Ayalur Kumar, Debashish Paul, Suman Majumder, Snehasis Mukherjee, Tatini Rakshit

## Abstract

Centromeric heterochromatin is maintained by a self-reinforcing loop between DNA methylation (5mC) and histone H3 lysine 9 trimethylation (H3K9me3), yet whether this coupling is preserved at the level of individual chromosomes in cancer has been inaccessible to conventional epigenomic assays, which average signals across cell populations and cannot resolve single-chromosome heterogeneity. Here, we use antibody-functionalized atomic force spectroscopy to simultaneously map 5mC and H3K9me3 at the centromeres of individual metaphase chromosomes from non-tumorigenic, luminal, and triple-negative breast cancer (TNBC) cell models. Co-localized 5mC/H3K9me3 sites are mechanically stiffer than sites carrying either mark alone across the panel, revealing a cooperative biophysical signature of epigenetic crosstalk. Luminal chromosomes acquire the highest mark density, co-localization, and stiffness, whereas TNBC chromosomes show reduced density and diminished, though not absent, co-localization and stiffness, converging toward the low baseline of non-tumorigenic epithelium. TGF-β1-induced epithelial-mesenchymal transition reproduces this attenuated phenotype in luminal cells, linking it directly to a process central to TNBC aggressiveness. These biophysical parameters classify cancer versus non-tumorigenic chromosomes with near-perfect, cross-validated accuracy, and further resolve the two TNBC subtypes using colocalization-derived features alone (macro-F1 = 0.933 ± 0.026). These results establish the mechanical consequence of epigenetic mark co-localization as a single-chromosome-resolvable signature of centromeric organization, attenuated in aggressive breast cancer, and demonstrate a force-spectroscopy platform for mechano-epigenetic profiling of cancer chromosomes.

**TOC:** 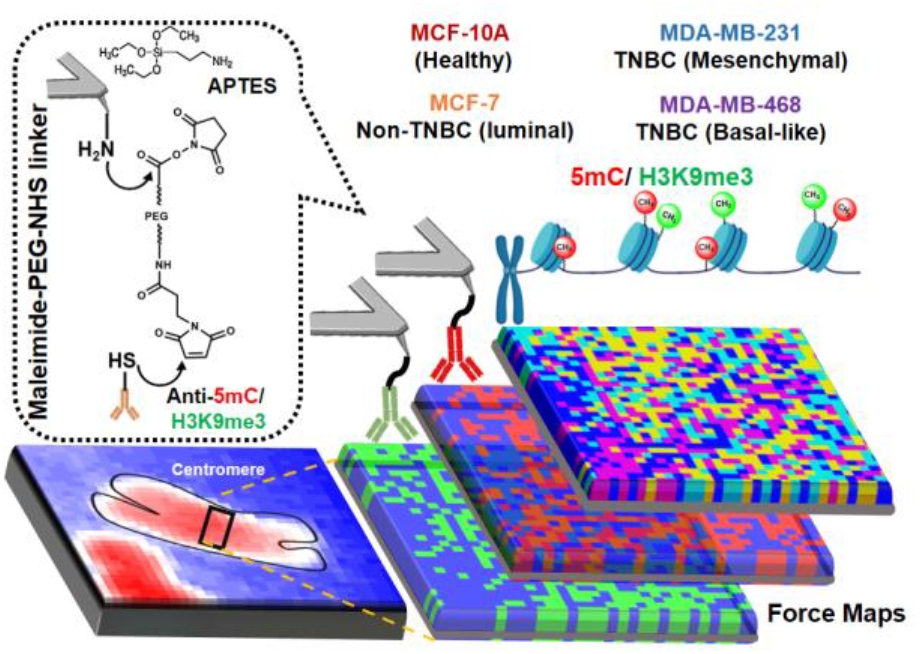

## Introduction

Breast cancer remains the most commonly diagnosed cancer in women worldwide, and triple-negative breast cancer (TNBC), lacking estrogen receptor, progesterone receptor, and HER2 expression, remains its most therapeutically intractable subtype, with treatment still largely confined to cytotoxic chemotherapy and persistently high rates of relapse^1^. A growing body of evidence implicates epigenetic dysregulation, alongside genetic mutation, as a driver of TNBC’s aggressive phenotype: DNA methylation (5-methylcytosine, 5mC) and histone H3 lysine 9 trimethylation (H3K9me3) are the two dominant repressive marks that establish and maintain constitutive heterochromatin, and at pericentromeric satellite repeats these marks are mechanistically coupled, 5mC recruits methyl-CpG-binding proteins that in turn deposit H3K9me3, and the resulting H3K9me3-HP1 complexes reinforce DNA methylation in a self-stabilizing loop^2–5^. This feedback loop establishes the boundaries of the CENP-A-containing core centromere, and its disruption is a recognized route to chromosomal instability (CIN), a hallmark of TNBC that drives tumor heterogeneity, metastatic capacity, and therapeutic resistance^2,5,6^.

Because 5mC and H3K9me3 physically coexist within a few nanometers of one another on the same nucleosome, their coordinated behavior should, in principle, be detectable as a local, physically coupled event with its own mechanical signature^7–9^. Yet every method currently used to study this relationship: whole-genome bisulfite sequencing, ChIP-seq, mass spectrometry-based histone profiling, and confocal immunofluorescence reports a population average across millions of cells and megabases of genome, collapsing exactly the spatial co-occurrence that defines the phenomenon^6,10,11^. Centromeric and pericentromeric DNA is additionally underrepresented in short-read sequencing, owing to its repetitive structure, compounding the problem^12^. Whether 5mC-H3K9me3 co-localization carries an independent mechanical signature, and whether the degree of co-localization differs systematically across breast cancer subtypes at single-chromosome resolution, has therefore remained inaccessible to direct observation^13^.

Single-molecule atomic force spectroscopy (AFS) offers a route around this limitation. AFM probes functionalized with epitope-specific antibodies can interrogate molecular recognition events with piconewton force resolution while simultaneously imaging the underlying structure, allowing force measurements to be spatially registered against real chromosome morphology^14^. This capability has previously been used to probe isolated protein-DNA and antibody-antigen interactions, but, to our knowledge, it has not been applied to co-map two distinct epigenetic marks on the same chromosome, nor to ask whether their spatial co-occurrence carries an independent mechanical signature^15^.

Here, we use dual-antibody AFS, anti-5mC, and anti-H3K9me3 probes applied to the same centromeric region of individual metaphase chromosomes to address three questions in a panel spanning non-tumorigenic epithelium (MCF-10A), luminal breast cancer (MCF-7), and two transcriptionally distinct TNBC subtypes (basal-like MDA-MB-468; mesenchymal-like MDA-MB-231). First, does 5mC-H3K9me3 co-localization carry a mechanical signature distinct from either mark alone? Second, does the degree of this co-localization differ systematically between luminal and triple-negative breast cancer? Third, is any such difference a fixed property of the cancer subtype, or can it be induced directly by epithelial-mesenchymal transition (EMT), the process most strongly linked to TNBC’s invasive and chemo-resistant phenotype?^16,17^ We complement the single-chromosome force measurements with bulk biochemical validation (ELISA, circular dichroism, FTIR) on the same cell panel. Five predefined supervised machine-learning experiments are then tested (i) cancer-versus-normal separation, (ii) three-class differentiation of MCF-7, MDA-MB-231, and MDA-MB-468 using morphology or non-colocalized molecular-mechanical features, and (iii) TNBC subtype differentiation using non-colocalized or colocalized representations.

**Scheme 1.**
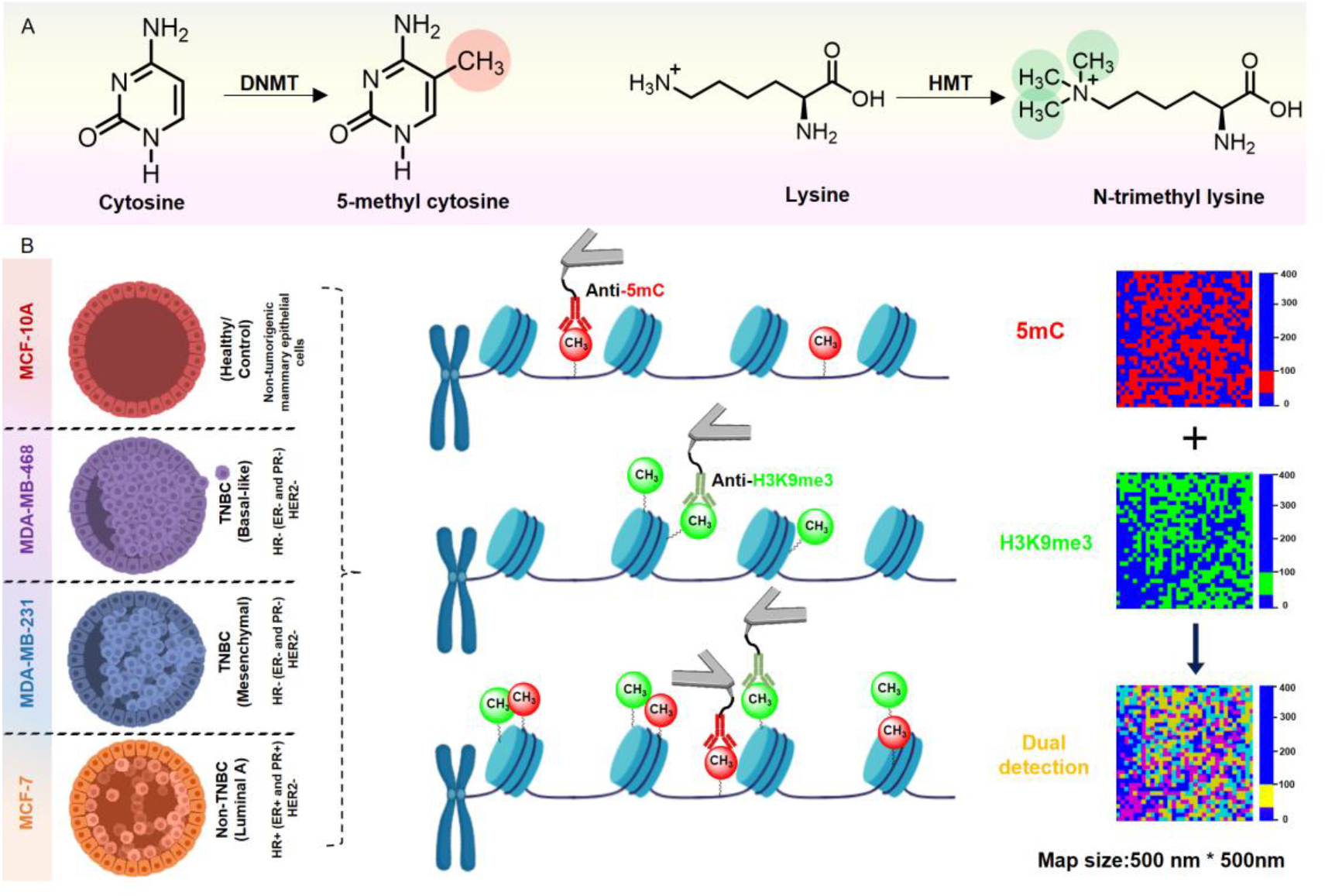
Schematic illustration of epigenetic modifications and AFM-based nanoscale mapping (A) Enzymatic methylation of cytosine to 5mC mediated by DNMT, and trimethylation of lysine to histone H3 lysine 9 (H3K9me3) mediated by HMT. (B) AFM-unbinding strategy employing anti-5mC and anti-H3K9me3 functionalized tips to probe methylated DNA and H3K9me3, respectively, on breast cell-derived chromosomes’ centromeres (MCF-7, MDA-MB-231, MDA-MB-468, and MCF-10A). Representative adhesion force maps (500 nm × 500 nm) illustrate the spatial distribution of methylated DNA and H3K9me3, and their nanoscale co-localization, highlighting regions of close proximity. (DNA methyltransferase: DNMT, histone methyltransferase: HMT, 5mC: 5-methylcytosine, H3K9me3: N-trimethyl lysine/ Histone H3 Lysine 9 Trimethylation).

## Results

Bulk epigenetic and structural differences distinguish the breast cancer panel. We first established that the four cell models, non-tumorigenic MCF-10A, luminal MCF-7, and the TNBC lines MDA-MB-468 (basal-like) and MDA-MB-231 (mesenchymal-like), carry distinct global epigenetic states before turning to single-chromosome measurements. Global DNA methylation, quantified by 5mC ELISA, was highest in MCF-7 (15.3%), intermediate in MDA-MB-231 (5.4%) and MDA-MB-468 (4.3%), and lowest in MCF-10A (1.2%); global H3K9me3 followed the same rank order (12.5%, 6.7%, 8.2%, and 3.1%, respectively; **Fig. 1A**). Circular dichroism of isolated histones showed higher α-helical and lower β-sheet content in MCF-10A and MDA-MB-231 relative to MCF-7 and MDA-MB-468 (**Fig. 1B, Table S1),** consistent with methylation-associated differences in histone secondary structure^18,19^. ATR-FTIR of isolated genomic DNA showed higher integrated absorbance (area under curve, AUC) in the 2850–3000 cm⁻¹ CH₂/CH₃ stretching region, diagnostic of the methyl moiety of 5-methylcytosine in cancer-derived DNA relative to MCF-10A (AUC = 0.80, 0.69, 0.61, and 0.52 for MCF-7, MDA-MB-231, MDA-MB-468, and MCF-10A, respectively, a 1.1–1.5-fold enrichment over MCF-10A; **Fig. 1C, D**, **Table S3**; full peak assignments in **Table S2**), in the same rank order as the ELISA data^20^. Together, the ELISA and CH₂/CH₃ FTIR data establish that this panel spans a genuine gradient of global epigenetic states, with luminal cancer carrying the highest bulk 5mC/H3K9me3 burden of the four lines and non-tumorigenic epithelium the lowest, a ranking that reframes the question addressed in the remainder of this study: what happens to the coordination between these two marks, at single-chromosome resolution, as cells transition from this luminal-high state to the triple-negative phenotype?

**Figure 1.**
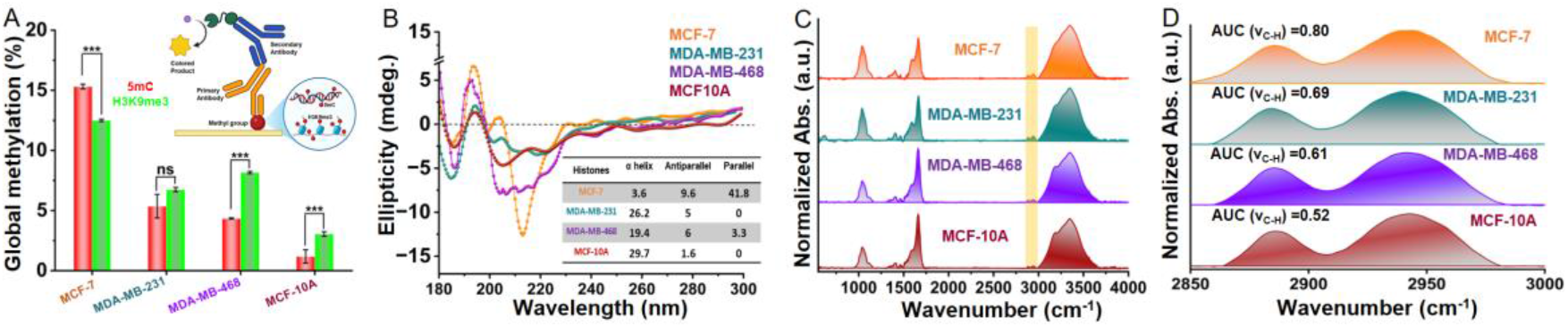
Bulk characterization of isolated genomic DNA and histones. (A) Quantification of global DNA methylation (5-methylcytosine, 5mC) and histone modification levels (H3K9me3) across all cell-derived samples. (B) Circular dichroism (CD) spectra of isolated histones from different cell types, with the inset table summarizing the estimated secondary structure composition (%). (C) FT-IR average spectral profile and (D) Integrated area under the curve (AUC) for genomic DNA, corresponding to the DNA methylation-associated -CH₃ stretching vibrations within the 2850–3000 cm⁻¹ spectral region.

### Dual-antibody force spectroscopy maps 5mC and H3K9me3 at individual centromeres

The panel spans the breast cancer receptor subtypes: MCF-10A (non-tumorigenic, HR-/HER2-mammary epithelium), MCF-7 (non-TNBC, luminal A, HR+/HER2-), and the two TNBC lines MDA-MB-231 (mesenchymal, HR-/HER2-) and MDA-MB-468 (basal-like, HR-/HER2-), each probed by dual-antibody AFS as illustrated in **Scheme 1**. To resolve the coupling question at the chromosome level, we performed high-resolution AFM imaging and antibody-based force spectroscopy (AFS) on metaphase chromosomes isolated from all four lines (**Fig. 2A–H**). Centromeric constrictions were clearly resolved in every chromosome, and quantitative morphometry revealed significant cell-line-dependent differences in centromeric height, width, and roughness (**Fig. 2I–K, Table S4**). Height was lowest in MCF-7 and increased progressively through MDA-MB-231 and MCF-10A to a maximum in MDA-MB-468; width was reduced in both TNBC lines relative to MCF-7, while MCF-10A did not differ significantly from MCF-7. Roughness, notably, did not track a simple cancer-versus-normal or luminal-versus-TNBC pattern: it increased monotonically from MCF-7 (lowest) through MDA-MB-231 and MDA-MB-468 to MCF-10A, which displayed the highest centromeric roughness of the panel, an exception to the otherwise cancer-associated morphological trend that we do not attempt to over-interpret here.

**Figure 2.**
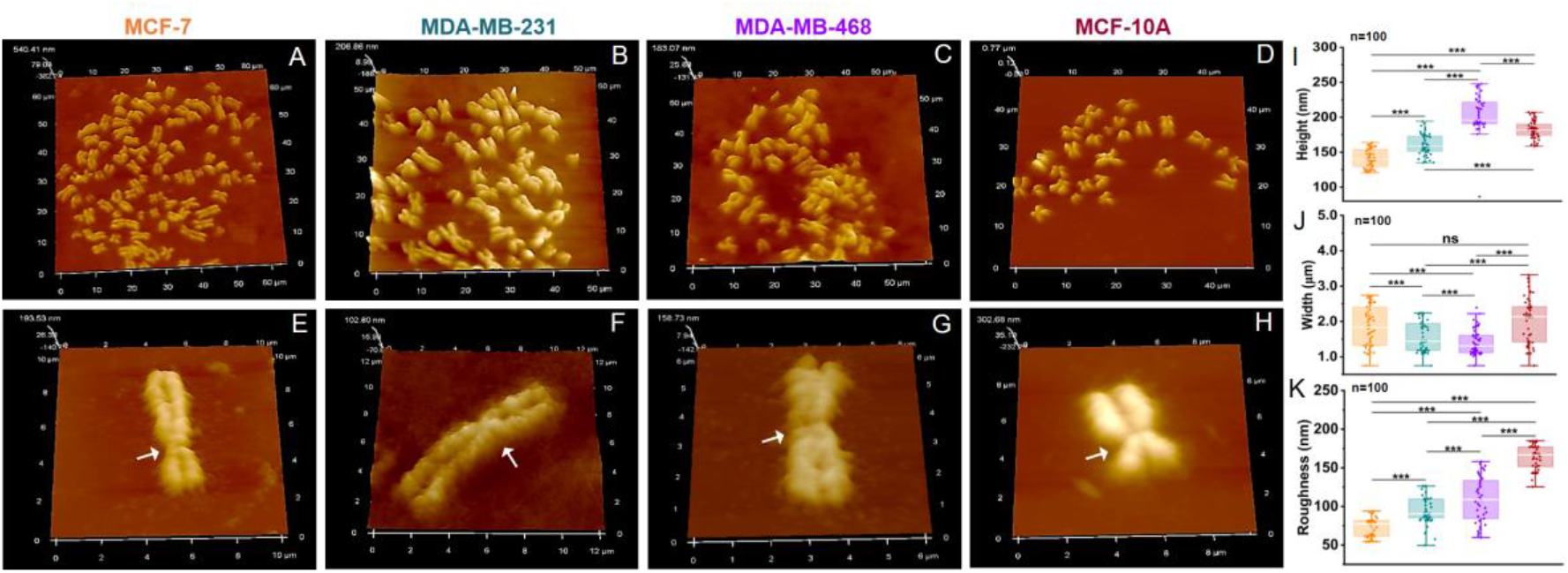
AFM-based 3D topographical analysis of chromosomes in physiological PBS buffer. (A, E) MCF-7, (B, F) MDA-MB-231, (C, G) MDA-MB-468, and (D, H) MCF-10A chromosomes showing representative 3D reconstructed images. Box plot quantification of centromeric regions depicting (I) height, (J) width, and (K) surface roughness (Ra) across all cell types. Statistical significance was evaluated using an unpaired t-test, with ns indicating non-significant differences.

We then functionalized silicon-nitride AFM tips with anti-5mC or anti-H3K9me3 antibodies via a flexible PEG spacer and recorded force-distance curves across a 500 × 500 nm² centromeric grid (32 × 32 pixels) for each chromosome, applying a 20–100 pN force window to isolate specific antibody-epitope unbinding events from non-specific adhesion. Individual retraction curves were classified as silent (no adhesion), mono-specific (a single rupture event), multi-specific (two or more sequential rupture events on the same curve, indicative of multivalent tip-chromosome engagement), or non-specific (a force signature inconsistent with antibody-epitope unbinding) (**Fig. S2**; representative curves for each chromosome in **Fig. S5**). The most probable unbinding force for both antibody systems were between 35–50 pN (**Fig. S6C–E**), and unbinding force scaled with the logarithm of loading rate as predicted by the Bell-Evans model (**Fig. S6A, B**), yielding a single dissociation barrier for each interaction: xβ = 0.88 ± 0.03 nm and koff = 1.54 ± 0.7 s⁻¹ for anti-5mC, and xβ = 0.37 ± 0.01 nm and koff = 0.56 ± 0.07 s⁻¹ for anti-H3K9me3 (**Table S5**). The corresponding activation energies (ΔG‡ = 17.1 and 17.78 kcal/mol, respectively) place the anti-H3K9me3 complex at a substantially longer bond lifetime (1/koff ≈ 1.8 s) than the anti-5mC complex (≈0.65 s, a 2.7-fold difference), indicating that the two recognition events, while comparable in peak force, sit in kinetically distinct regimes. Antibody-blocking controls reduced binding probability from 35.2% and 36.4% (5mC and H3K9me3, respectively) to 2.5% and 2.6% (Fig. S7), confirming that the measured events reflect specific epitope recognition rather than tip- or substrate-derived artifacts. Topography and force maps of the same 500 nm² region, acquired sequentially and co-registered by an in-house drift-correction routine (**Scheme S1**; example alignment in **Fig. S3D**), allowed each binding event to be assigned a precise position on the underlying chromosome structure^21^. This registration was reproducible across hierarchical scales, from repeated maps of the same centromere and large-area chromosome scans to paired single-antibody and merged maps on individual chromosomes and centromeric regions specifically (**Fig. S3**), and across five independent replicate maps per cell line (**Fig. S4A–D**).

### Centromeric mark density and co-localization are highest in luminal cancer, lowest in non-tumorigenic epithelium

Representative dual-recognition maps showed spatially clustered, specific binding for both 5mC and H3K9me3 at the centromere of every chromosome examined (**Fig. 3A**). Binding probability, the fraction of scanned pixels showing a specific unbinding event followed the same rank order as the bulk ELISA data: highest in MCF-7 (28.2 ± 2.1% for 5mC, 25.2 ± 2.2% for H3K9me3), intermediate in MDA-MB-231 (18.1 ± 4.2%, 12.9 ± 2.0%) and MDA-MB-468 (11.7 ± 1.1%, 18.7 ± 1.2%), and lowest in MCF-10A (2.1 ± 0.6%, 3.7 ± 0.4%) (**Fig. 3B, Table S4**). MDA-MB-468 was the sole exception to an otherwise consistent 5mC-over-H3K9me3 ordering within each line: its H3K9me3 binding probability exceeded its own 5mC value, a reversal not seen in MCF-7, MDA-MB-231, or MCF-10A, and one we do not have a mechanistic explanation for from the present data. The probability of finding a co-localized (“merged”) 5mC/H3K9me3 signal at the same pixel followed the same overall rank order (**Fig. 3C**): 5.7 ± 0.3% (MCF-7), 2.1 ± 0.6% (MDA-MB-231), 1.5 ± 0.3% (MDA-MB-468), and 0.4 ± 0.08% (MCF-10A). This single-chromosome measurement therefore recovers, at nanometer resolution, the same luminal-high/epithelial-low gradient established by bulk ELISA, and extends it with information the bulk assay cannot provide: TNBC chromosomes retain a measurable, non-zero degree of 5mC-H3K9me3 co-localization, intermediate between the luminal-high and epithelial-low extremes rather than showing a complete loss of coupling. We return to the significance of this partial, graded reduction, rather than an outright loss, in the Discussion.

**Figure 3.**
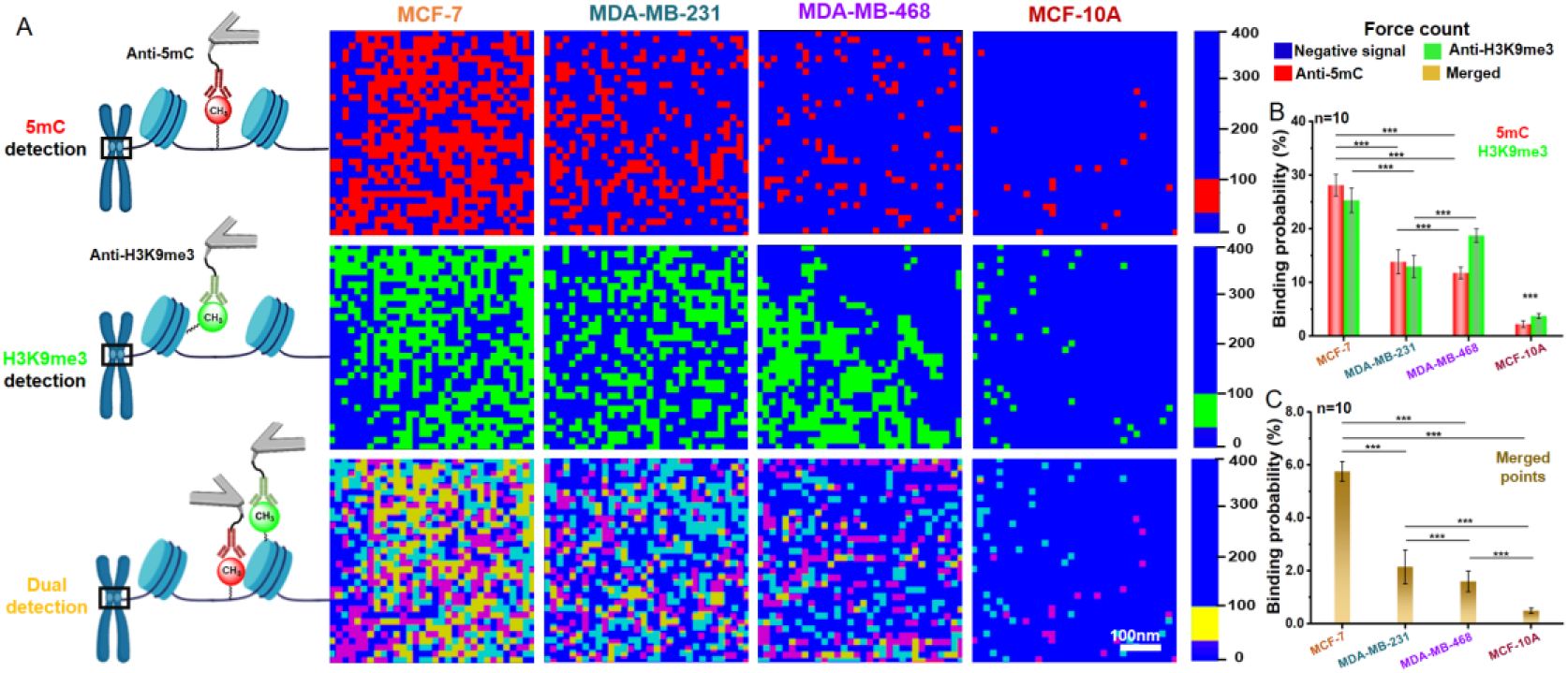
AFM-based force mapping reveals the spatial distribution and co-localization of DNA and histone methylation marks on single chromosomes. (A) High-resolution Force maps (500*500nm^2^;32*32 pixels) acquired from the centromeric regions of chromosomes isolated from MCF-7, MDA-MB-231, MDA-MB-468, and MCF-10A cells. The schematic (right) illustrates the strategy employed for single- and dual-antibody detection during AFM unbinding measurements. Percentages of Binding probabilities occupied by the positive pixel (B) for 5mC and H3K9me3, and (C) for merged points of all chromosomes.

### Co-localized 5mC/H3K9me3 sites are mechanically stiffer than either mark alone

We next asked whether the spatial co-occurrence of 5mC and H3K9me3 carries an independent mechanical signature. Retraction force curves showing specific adhesion events, were fitted with the worm-like chain (WLC) model to extract contour length (Lc) and, from the stretching force and unbinding distance, a stretching modulus (S) for each interaction (**Fig. 4A**). A subset of curves showed two or more sequential WLC-fittable rupture events rather than one (**Fig. 4B**), consistent with the multi-specific curve category defined in **Fig. S2** and providing direct evidence that a subset of tip-chromosome engagements involves more than a single antibody-epitope pair a plausible physical basis for the cooperative mechanical effect described below. For single-mark interactions, Lc was shortest in MCF-7 for both 5mC and H3K9me3 and longest in MDA-MB-468 (5mC) and MDA-MB-231 (H3K9me3) (**Fig. 4C, D**), and correspondingly, S was highest in MCF-7 for both marks (124.6 ± 6 pN for 5mC; 130.7 ± 17 pN for H3K9me3) and lower in the two TNBC lines (5mC: 90.1 ± 23 pN in MDA-MB-231, 102 ± 7 pN in MDA-MB-468; H3K9me3: 110.3 ± 5 pN in MDA-MB-231, 128.7 ± 18 pN in MDA-MB-468; **Fig. 4G, H**). MDA-MB-231 and MDA-MB-468 chromosomes also showed larger unbinding distances than MCF-7 (**Fig. S8C, D**), together indicating a more extensible, mechanically compliant chromatin fiber at single-mark sites in the TNBC lines.

**Figure 4.**
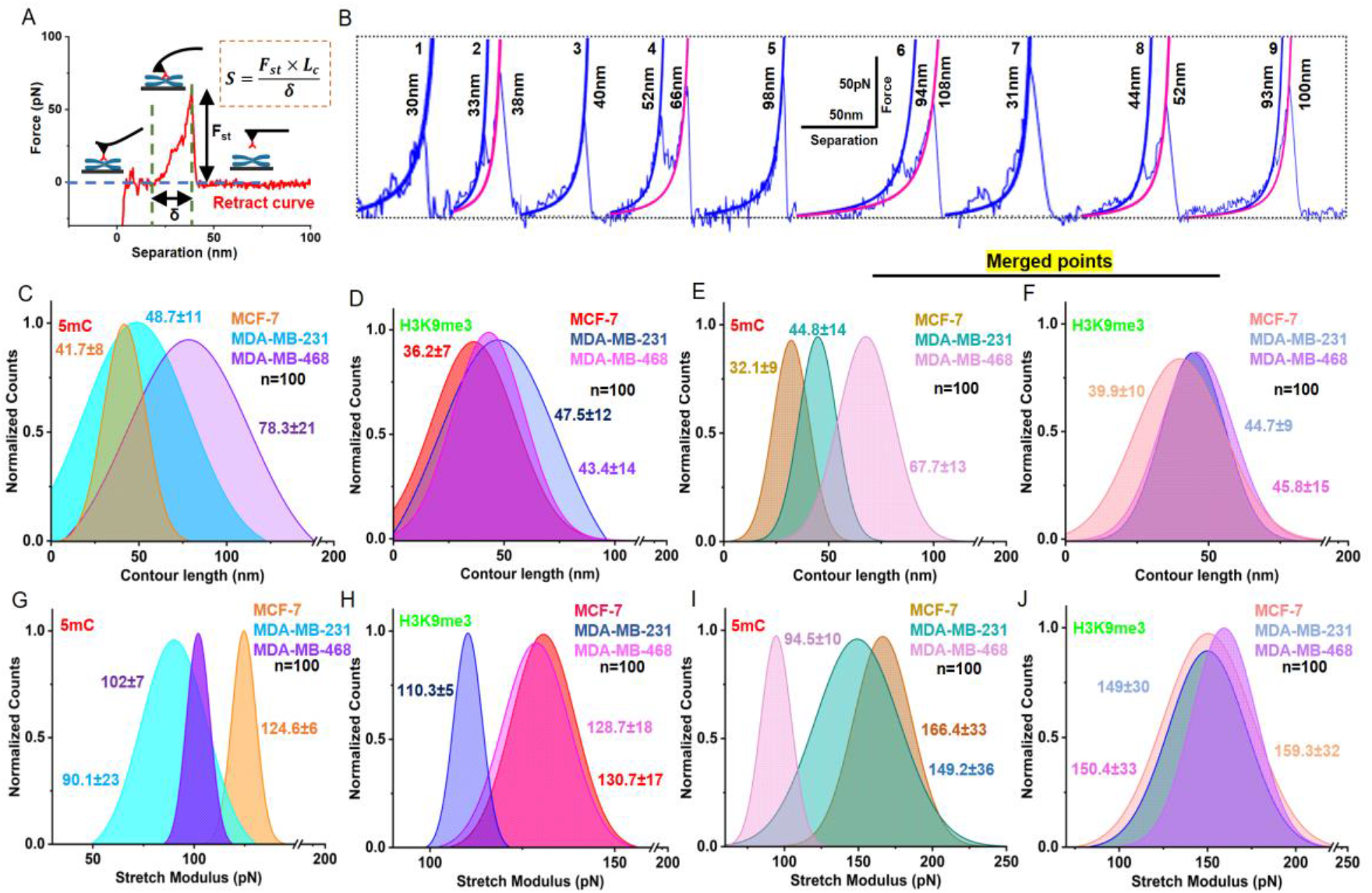
Force spectroscopy of chromatin via stretching modulus analysis. (A) Representative retract curve showing chromatin stretching (Fst=stretching force); the dotted blue line represents the baseline. (B) Sample retract curves for single and multiple rupture events for all cases; corresponding LCs were quantified by WLC fitting. Normalized Gaussian-fitted distribution plots of contour lengths for (C) 5mC and (D) H3K9me3 positive binding curves of all centromeres. Distribution plot of contour lengths for (E) 5mC and (F) H3K9me3 positive binding curves, where dual detection (merged points) of all chromosomes was observed, and the mean LC ± SD was quantified. Corresponding stretching modulus (S) values were obtained for (G) 5mC and (H) H3K9me3, and for (I) 5mC and (J) H3K9me3, where dual detection was observed (merged points) across all centromeres. The numbers indicate the mean ± standard error obtained from the Gaussian fits. Stretching modulus was calculated using S = Fst x LC/δ from force-separation curves, where δ is the unbinding distance, and LC is the contour length, n=100.

At sites where 5mC and H3K9me3 were co-localized, stretching modulus was higher than at single-mark sites in five of the six mark/cell-line combinations examined: for 5mC, merged-site S increased to 166.4 ± 33 pN in MCF-7 (from 124.6 ± 6 pN single-mark) and 149.2 ± 36 pN in MDA-MB-231 (from 90.1 ± 23 pN); for H3K9me3, merged-site S increased to 159.3 ± 32 pN in MCF-7, 149 ± 30 pN in MDA-MB-231, and 150.4 ± 33 pN in MDA-MB-468 (from single-mark values of 130.7 ± 17, 110.3 ± 5, and 128.7 ± 18 pN, respectively; **Fig. 4I, J**; statistics in **Fig. S8**). The sole exception was 5mC at merged sites in MDA-MB-468, where modulus (94.5 ± 10 pN) was statistically indistinguishable from the single-mark value (102 ± 7 pN) rather than elevated. Two-dimensional kernel density estimation of adhesion force versus unbinding distance for the individual and merged interactions (**Fig. S9**) corroborated the overall pattern of reduced mechanical rigidity in TNBC: MCF-7 occupied a comparatively concentrated force-distance regime for both marks, whereas MDA-MB-468 in particular showed a markedly broader distribution extending to much larger unbinding distances, consistent with the more extensible, less rigid chromatin fiber inferred from the WLC fits above (MCF-10A was not included in this analysis). Taken together, these data indicate that co-occurrence of 5mC and H3K9me3 at the same centromeric locus is associated with a *cooperative,* mechanically stiffening effect^22^ that is largely conserved across cell lines and is attenuated, though not completely absent, in TNBC chromosomes, the biophysical signature that gives this study its central claim.

### Publicly available data place EMT-associated genes at variable centromeric distances, accompanied by cell-line-specific DNA methylation of EMT genes

To investigate whether centromeric positioning correlates with the regulatory status of EMT-associated genes, we first evaluated publicly available datasets. Using the Hallmark epithelial-mesenchymal transition gene set, we computed the genomic distance of each EMT-associated gene from the centromere of its resident chromosome, expressed as a signed distance (upstream/downstream) in megabases; EMT genes were distributed across essentially every chromosome at variable centromeric distances, with no single chromosome or arm disproportionately enriched (**Fig. S1A**). We then examined DNA methylation of these same EMT-associated genes in MDA-MB-231, MCF-7, MDA-MB-468, and MCF-10A using publicly available methylation array data (GSE42944), expressed as Z-scored average methylation per gene (**Fig. S1B**)^23–26^ . This analysis revealed clear, cell-line-dependent differences in EMT gene methylation across the panel, consistent with the existence of a distinct, cell-line-specific epigenetic regulatory state for EMT genes independent of our own AFM measurements, supporting the plausibility of a mechanistic link between global/centromeric methylation state and EMT gene regulation that we test directly, at the single-chromosome level, in the induction experiment below. We note that a directly comparable, adequately powered public H3K9me3 dataset covering all four lines with sufficient centromeric/pericentromeric resolution was not available, so this genomic-context analysis was restricted to DNA methylation; the H3K9me3 side of our central finding rests entirely on the AFM-based measurements described above, a point we return to in the Discussion.

### TGF-β1-induced EMT reproduces the TNBC methylation-coupling phenotype in luminal cells

To test whether the reduced density and co-localization seen in TNBC chromosomes could be induced directly, rather than reflecting a fixed subtype-intrinsic state, we treated MCF-7 cells with TGF-β1 (10 ng mL⁻¹) to induce EMT and isolated metaphase chromosomes for AFS (**Fig. S10A–C**). TGF-β1 treatment significantly reduced binding probability for both 5mC and H3K9me3 individually (**Fig. S10F**) and reduced the probability of co-localized recognition events (**Fig. S10G**). Contour length increased and stretching modulus decreased for both marks following treatment (**Fig. S10H, I**), mirroring the more compliant, less coupled phenotype of native TNBC chromosomes. Principal-component analysis of the full AFM-derived parameter set placed TGF-β1-treated MCF-7 chromosomes in the same region of PC space as MDA-MB-468 (basal-like TNBC), clearly displaced from untreated MCF-7 and from the distinct MDA-MB-231 cluster (**Fig. S10J**), indicating that EMT induction alone is sufficient to drive a luminal chromosome toward a specifically basal-like TNBC centromeric methylation-coupling state.

### Machine learning resolves cell-line identity and cancer subtype from single-chromosome biophysical fingerprints

Finally, we asked whether the combination of morphological, density, colocalization, and mechanical parameters measured on a single chromosome is sufficient to classify cell-of-origin and cancer subtype. Using an 11-feature set (centromere height, width, and roughness; 5mC and H3K9me3 binding probability; contour length, stretching modulus, and unbinding distance for both marks), we evaluated five classification experiments — cancer versus normal, three-class morphology, three-class non-colocalized, TNBC non-colocalized, and TNBC colocalized (**Fig. 5A–E**) — across four classifiers (random forest, SVM, Gaussian naive Bayes, and weighted k-NN), assessed by repeated nested cross-validation (10 repetitions; 5 stratified outer folds; 4 inner folds for hyperparameter selection; **Fig. S11**; **Tables S6, S7**). For cancer-versus-normal classification using all 11 features, random forest achieved accuracy, balanced accuracy, macro-F1, macro-AUC, and MCC all equal to 1.000, while SVM and Gaussian naive Bayes achieved macro-F1 of 0.978 ± 0.012 and 0.989 ± 0.000, respectively (**Fig. S12**)^27–29^. Morphology alone was substantially weaker for the harder three-class (MCF-7/MDA-MB-231/MDA-MB-468) discrimination (best macro-F1 = 0.723 ± 0.026, Gaussian naive Bayes); the confusion was concentrated specifically between MCF-7 and MDA-MB-231, each misclassified as the other in 37% of instances, while MDA-MB-468 was already well separated by morphology alone (91% correct). Replacing morphological features with the non-colocalized molecular-mechanical feature set improved macro-F1 for every classifier (by 0.084–0.196, **Fig. S13**) and resolved most of this specific confusion: random forest achieved the best three-class performance (accuracy 0.901 ± 0.014, macro-F1 0.901 ± 0.014, macro-AUC 0.967 ± 0.004), with class-wise recall of 0.947 for MCF-7, 0.847 for MDA-MB-231, and 0.910 for MDA-MB-468. For TNBC subtype separation specifically, random forest using non-colocalized features achieved macro-F1 = 0.925 ± 0.018 (balanced across subtypes: MDA-MB-231 F1 = 0.926, MDA-MB-468 F1 = 0.924), while SVM using the seven-feature colocalized representation achieved the overall best subtype separation (macro-F1 = 0.933 ± 0.026, macro-AUC = 0.981 ± 0.011), with random forest comparable (macro-F1 = 0.932 ± 0.012). The benefit of colocalization was classifier-dependent: relative to the non-colocalized representation, macro-F1 increased by 0.077 for SVM and 0.007 for random forest, but decreased by 0.039 for weighted k-NN and 0.025 for Gaussian naive Bayes (**Fig. S14**), indicating that colocalized epigenetic-mechanical features are informative but not universally superior across decision architectures. Together, these results show that single- hromosome AFS parameters, without any sequence-level information, are sufficient to separate normal, luminal, and TNBC chromosomes, resolving specifically the luminal/mesenchymal-TNBC confusion that morphology alone cannot, and to resolve the two TNBC subtypes from one another using the same colocalization-derived features that carry the study’s central mechanistic finding (**Fig. 5; Tables S8–S11**). These results establish strong internal cross-validation separability of these cell-line chromosome measurements, rather than demonstrating clinical diagnostic performance.

**Figure 5.**
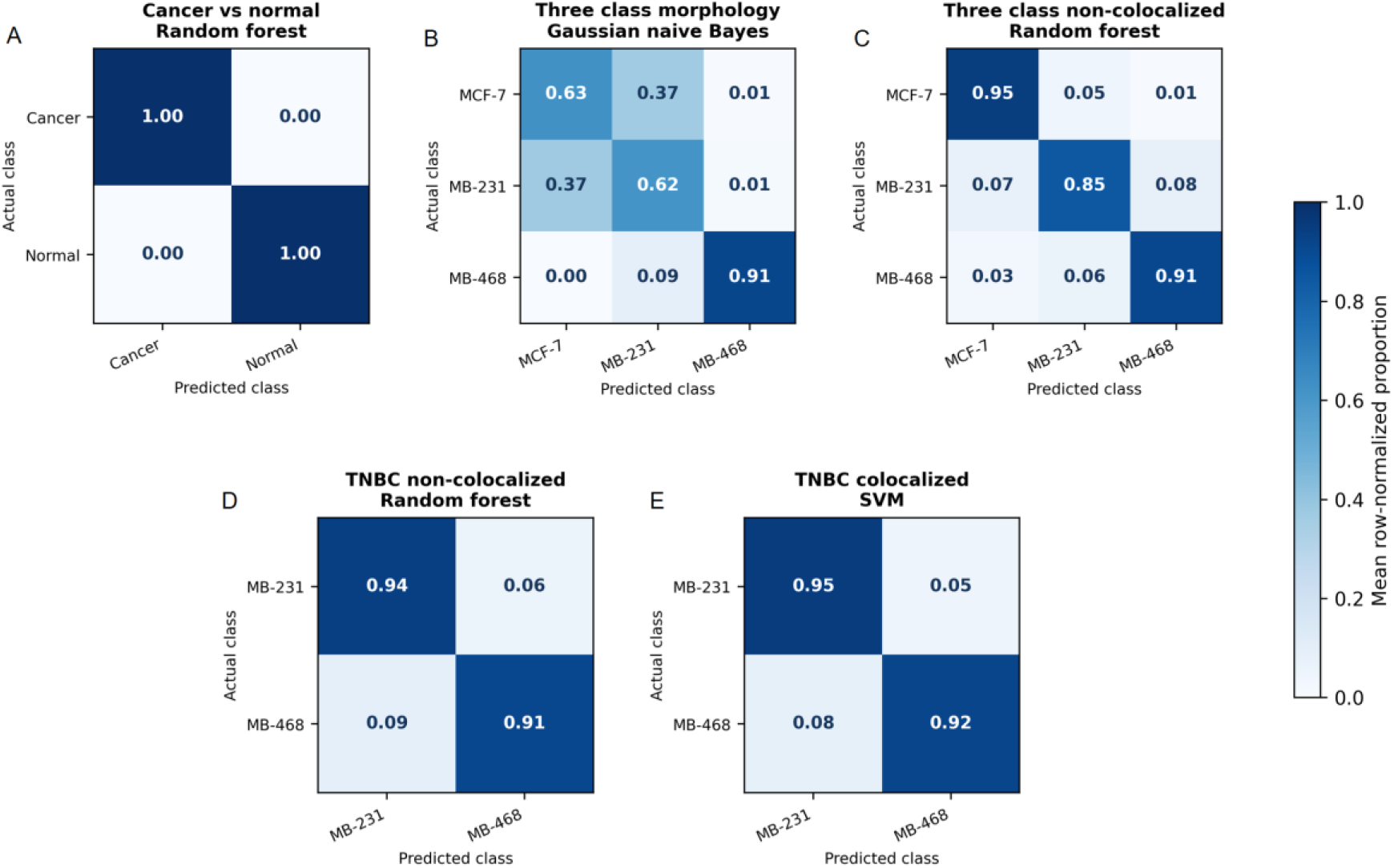
Machine-learning analysis of AFM derived parameters across chromosomes. Average row-normalized confusion matrix (%) for the best-performing pipeline across (A) Cancer and normal chromosomes with all AFM parameters (B) Non-TNBC and TNBC chromosomes with morphological features (Height, Width and Roughness) (C) Non-TNBC and TNBC chromosomes with force spectroscopy features (Unbinding Distance, Binding Probability, Contour Length and Stretch Modulus) (D) TNBC subtypes with Unbinding Distance, Binding Probability, Contour Length and Stretch Modulus features (E) TNBC subtypes with Unbinding Distance, Binding Probability, Contour Length and Stretch Modulus features for the colocalized/merged dataset. Diagonal elements represent correctly classified samples, while off-diagonal elements indicate misclassifications between classes. SVM: support vector machines

## Discussion

Ensemble epigenomic methods have established that 5mC and H3K9me3 are mechanistically coupled at pericentromeric heterochromatin and that this coupling is disrupted in cancer, but because these methods average across millions of cells and megabases of genome, they cannot say whether coupling is lost gradually, uniformly, or heterogeneously at the level of a single chromosome, nor whether the physical co-occurrence of the two marks has any mechanical consequence distinct from their individual presence^22,30–32^. The single-chromosome force spectroscopy presented here addresses both gaps directly, and the central finding is a mechanical one: wherever 5mC and H3K9me3 co-occur at the same centromeric locus, the chromatin fiber is measurably stiffer than at sites carrying either mark alone, in five of six mark/cell-line combinations tested. This cooperative stiffening is, to our knowledge, the first direct biophysical demonstration that the well-established biochemical crosstalk between DNA and histone methylation has a corresponding mechanical signature at the single-locus level, consistent with a model in which co-occurring marks nucleate more extensive local chromatin compaction than either mark achieves alone^33^.

Layered onto this general rule, the four-line panel resolves a specific disease-relevant gradient, summarized schematically in **Scheme 2**. Luminal (MCF-7) chromosomes carry the highest density of both marks, the highest degree of co-localization (5.7 ± 0.3%), and the stiffest co-localized sites of the panel: a hypermethylated, tightly coupled, mechanically rigid centromeric state that we interpret as dense heterochromatin. Co-localization decreases progressively through MDA-MB-231 (2.1 ± 0.6%, “partially remodeled” chromatin) and MDA-MB-468 (1.5 ± 0.3%, “further remodeled”), reaching a minimum in non-tumorigenic MCF-10A (0.4 ± 0.08%, the most relaxed/decondensed state of the panel). TNBC chromosomes, in both the basal-like and mesenchymal-like subtypes, show reduced mark density, reduced co-localization probability, and lower stretching modulus at both single-mark and (in most cases) merged sites, converging toward, though not below, this low epithelial baseline. We deliberately avoid describing this as a loss or decoupling of the 5mC-H3K9me3 relationship: co-localized events remain clearly detectable in both TNBC lines (**Fig. 3C**), and the correct description is a graded attenuation of coupling and its mechanical consequence, superimposed on a cancer-associated hypermethylated state that luminal cells acquire and TNBC cells only partially retain. This distinction matters mechanistically: a chromosome that has lost coupling entirely and one that retains a diminished, still-cooperative coupling are, in principle, governed by different regulatory logic, and only single-chromosome measurement can distinguish them^5,30^.

The TGF-β1 induction experiment provides a causal, rather than purely correlative, link between this phenotype and EMT: driving luminal MCF-7 cells through EMT is sufficient to reduce mark density, reduce co-localization, and soften centromeric chromatin, reproducing the TNBC-like signature within a single, genetically matched cell background. Because EMT is itself a recognized driver of chromosomal instability in TNBC, this result places centromeric methylation-coupling attenuation as a plausible intermediate step between EMT induction and the compromised centromere function, impaired kinetochore assembly, weakened chromatid cohesion, and aneuploidy documented in aggressive breast cancer^34,35^. The machine-learning results extend the mechanistic observations to a quantitative classification setting: the complete 11-feature representation separated cancer from non-tumorigenic chromosome measurements under internal nested cross-validation, the non-colocalized molecular-mechanical representation improved three-class cell-line differentiation over morphology alone, and the colocalized representation supported high MDA-MB-231-versus-MDA-MB-468 discrimination. These complementary results show that the measured signature is biologically informative, though they do not yet establish diagnostic generalization beyond the cell lines studied.

**Scheme 2.**
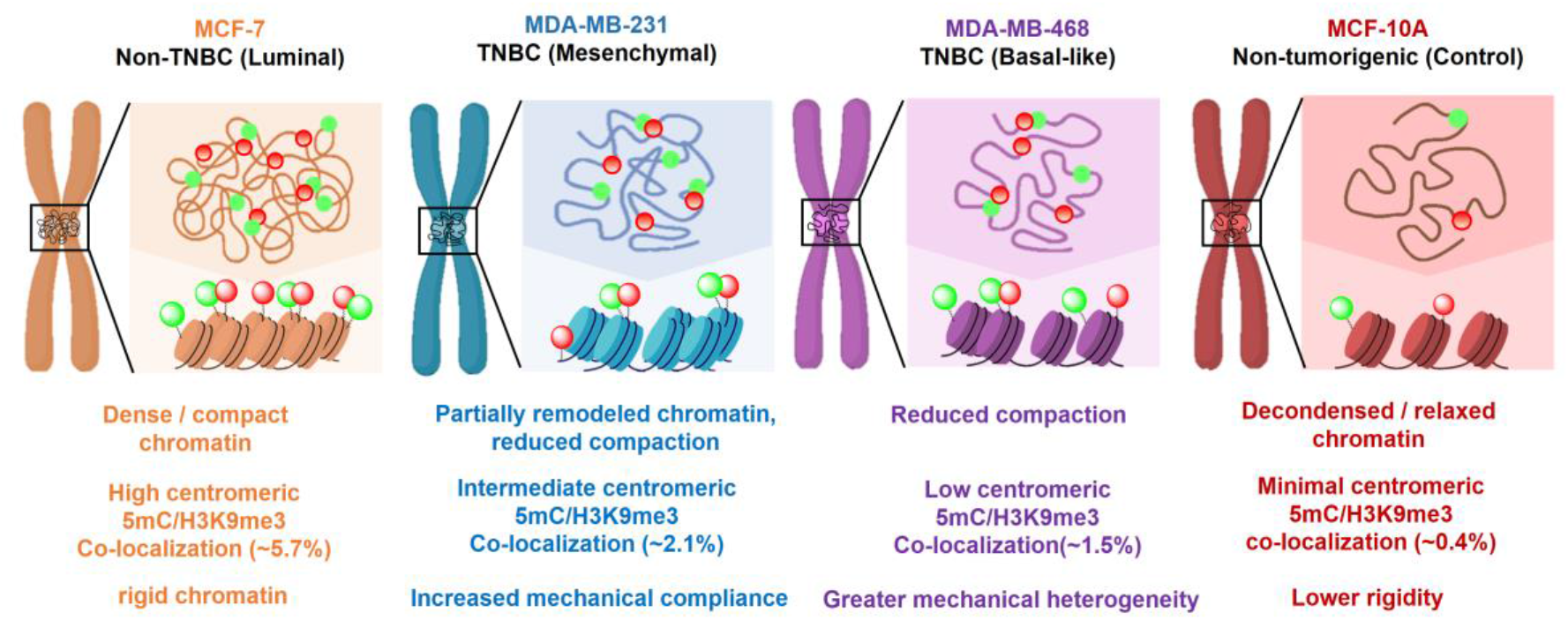
Proposed model of chromatin organization across breast cancer subtypes.

### Limitations bound the scope of these conclusions

AFS measurements on isolated, immobilized metaphase chromosomes may not exhibit native chromatin behavior in intact, living cells. The probe and linker chemistry (antibody tethering efficiency, PEG length heterogeneity) contributes to variability that we minimized but did not eliminate through blocking controls and force-threshold filtering. Distinguishing genuinely single-molecule unbinding events from closely spaced multiple interactions remains an intrinsic limitation of antibody-based AFS, and could inflate estimates of interaction density at the highest-probability sites^36^. The genomic-context analysis of EMT-associated gene methylation (**Fig. S1**) relied on public DNA methylation data alone; no publicly available H3K9me3 dataset offered sufficiently consistent coverage and centromeric/pericentromeric resolution across all four lines to support a comparable analysis for that mark, so the H3K9me3 component of our central finding rests on AFM-based measurements rather than on independent genomic corroboration. High-depth, centromere-resolved H3K9me3 profiling of all four lines, ideally spanning EMT induction, would meaningfully strengthen the genomic grounding of the biophysical phenotype reported here. The machine-learning analysis used repeated nested cross-validation to separate hyperparameter selection from outer-fold performance estimation and included 30 chromosome-level observations per cell line. Individual chromosome measurements were treated as the analysis units. Because chromosomes originating from a common culture, preparation, or acquisition batch may share batch-specific characteristics, stratified chromosome-level folds cannot rule out all sources of biological or experimental dependence unless the preparation identifiers are incorporated as grouping variables. The reported values should therefore be interpreted as internal proof-of-concept evidence of feature informativeness, not as independently validated diagnostic accuracy; confirmation using independently prepared samples and patient-derived material is required^37^.

These limitations notwithstanding, this work establishes single-molecule AFS as a platform capable of resolving a property, the mechanical consequence of epigenetic mark co-localization, that is invisible to every ensemble method currently used to study centromeric chromatin. Beyond breast cancer, the same dual-antibody force-mapping strategy is directly applicable to any system in which two chromatin marks are conjectured to interact mechanically rather than merely co-occur statistically, and the demonstration that a purely biophysical, sequence-agnostic parameter set can classify cancer subtype from single chromosomes suggests a route toward mechanoepigenetic diagnostics that complements, rather than replaces, existing genomic and transcriptomic classification schemes^29^.

Taken together, this study shows that centromeric 5mC and H3K9me3 are not merely co-located but mechanically cooperative. This cooperativity is present across non-tumorigenic, luminal, and triple-negative breast chromosomes but is quantitatively attenuated along a luminal-to-basal-to-mesenchymal-to-epithelial gradient, which EMT induction alone reproduces in luminal cells, and the resulting biophysical fingerprint is sufficient to classify chromosome identity without any sequence-level information. Single-chromosome force spectroscopy, therefore, adds a mechanical dimension, inaccessible to ensemble genomic and epigenomic assays, to the study of centromeric chromatin organization in cancer, and provides a template for asking the same co-localization/mechanics question in other cancers and other chromatin-mark pairs.

## Methods

Cell culture and metaphase chromosome isolation. MCF-7 (luminal A), MDA-MB-231 (basal-B TNBC), and MDA-MB-468 (basal-A TNBC) cells (NCCS, Pune, India) were cultured in high-glucose DMEM (Gibco) with 10% FBS (Gibco) and 1% penicillin/streptomycin (Gibco). MCF-10A (ATCC, USA) was cultured in DMEM/F12 supplemented with 5% horse serum (Himedia), 1% penicillin/streptomycin, 0.5 μg mL⁻¹ hydrocortisone (SRL), 100 ng mL⁻¹ cholera toxin (Sigma-Aldrich), and 10 μg mL⁻¹ insulin (SRL). Cells were maintained at 37 °C, 5% CO₂. Metaphase chromosomes were isolated from ∼10⁶ cells per batch by colcemid arrest (0.1 μg mL⁻¹, overnight), hypotonic swelling (0.075 M KCl, 37 °C, 20 min), and fixation (methanol: acetic acid, 3:1 v/v). Fixed chromosomes were treated with RNase (Sigma-Aldrich, R6513) and protease (Sigma-Aldrich, P6887), quantified by NanoDrop, and stored in fixative at 4 °C for up to 24 h. Only well-condensed metaphase chromosomes with paired sister chromatids and a clearly defined centromere were used; fragmented, overlapping, or ambiguous chromosomes were excluded uniformly across groups. Experiments used cells within passages 3-5 after thawing.

### Genomic DNA and histone extraction

Genomic DNA was extracted using the DNeasy Blood and Tissue Kit (Qiagen), quantified by NanoDrop, and assessed for purity by A260/A280 (yield range 50-100 ng μL⁻¹). Histones were extracted by nuclear isolation followed by acid extraction (0.5 N HCl, 10% glycerol), dialyzed against chilled acetic acid, air-dried, resuspended in nuclease-free water, and quantified by A280 (yield range 0.8-2.5 μg μL⁻¹); samples were stored at −20 °C.

### ELISA quantification of global 5mC and H3K9me3

Global DNA methylation was quantified with the MethylFlash Global DNA Methylation (5mC) ELISA Easy Kit (EpigenTek) using 100 ng input DNA per well, with absorbance read at 450 nm (BioTek Synergy H1) and quantity calculated from a standard curve as %5mC = (sample OD − negative OD)/(slope × S) × 100, where S is input DNA mass (ng). Global H3K9me3 was quantified with the EpiQuik Global Tri-Methyl Histone H3-K9 Quantification ELISA Easy Kit (EpigenTek) using an analogous protocol and calculation.

### Circular dichroism

CD spectra of extracted histones were recorded on a Jasco J-1500 spectrophotometer (25 °C, 180-300 nm, triplicate, averaged) and deconvoluted into secondary-structure content using CDNN software, cross-validated against the BeStSel server.

### ATR-FTIR spectroscopy

ATR-FTIR spectra of isolated genomic DNA (2 μL per sample) were recorded on a Nicolet iS20 spectrometer (Thermo Fisher) with a DTGS detector (500-4000 cm⁻¹, 4 cm⁻¹ resolution, 64-scan average, triplicate). Spectra were smoothed (Savitzky–Golay, 15-point window) and baseline-corrected by the second-derivative method.

### AFM imaging

Chromosome suspensions (50 μL) were deposited on freshly cleaned glass, incubated for 20 min, and washed with PBS. Imaging was performed in intermittent-contact mode in PBS (pH 7.4) on an Asylum MFP-3D (Asylum Research) using triangular silicon-nitride probes (SNL-10, spring constant 0.3 N m⁻¹, Bruker). Images were analyzed in Igor Pro; centromeric height, width, and roughness were measured on ∼50 chromosomes per cell line, and group differences were assessed by an unpaired t-test.

### AFM tip functionalization

SNL-10 probes were silanized with 5% (3-aminopropyl) triethoxysilane (APTES) in 5% ethanol/95% water, rinsed, and air-dried (15–30 min). Probes were then incubated with heterobifunctional MAL-PEG₂₄-NHS ester linker (2 mg mL⁻¹, ∼40 min), rinsed, and functionalized with anti-5-methylcytosine (Sigma-Aldrich, SAB2702243) or anti-H3K9me3 (Sigma-Aldrich, ZRB1888) antibody (10 μg mL⁻¹, 30 min, ambient), followed by a final Milli-Q rinse.

### AFM force spectroscopy

Force-distance curves and force maps were acquired in contact mode over a 500 × 500 nm² centromeric region per chromosome (32 × 32-pixel grid), with cantilever spring constant and deflection sensitivity calibrated before each session. Approximately 2,000-3,000 force curves were recorded across a single chromosome (∼10 chromosomes per cell line across the study), at a retraction speed of 500 nm s⁻¹ and applied force of 200-350 pN; for loading-rate-dependence measurements, retraction velocity was varied from 500-8,000 nm s⁻¹, and effective loading rate was calculated as the pre-rupture force-distance slope multiplied by retraction speed. Specific events were isolated from non-specific adhesion by a 20-100 pN force-threshold window. Topography was acquired first on the chromosome, zoomed to the centromere in a few steps, with an unmodified tip, followed immediately by force mapping with the antibody-functionalized tip at the same stage position, minimizing lateral drift; a closed-loop scanner-maintained tip positioning, and antibody-functionalized tips were periodically tested against bare glass to confirm the absence of non-specific adhesion. Blocking controls were pre-incubated with chromosomes with anti-5mC or anti-H3K9me3 antibody prior to AFS.

### Drift-corrected map registration

Consecutive AFM force maps acquired from the same centromeric region were registered to compensate for lateral drift before overlay, by applying basic affine transformations (translation and rotation) based on two reference points. A custom Python 3 code was used to determine the x-y displacement (affine transformation) between successive maps. The geometric centroid of the chromosome was extracted from the corresponding height distribution and used as the positional reference point for the affine transformation. One map was fixed while the subsequent map was translated pixel-by-pixel along the x and y axes, and the displacement (transformation) yielding the minimum centroid mismatch was selected as the drift correction, based upon the translation and rotation of the two chosen reference points. The corrected maps were then overlaid sequentially to spatially compare molecular binding features. This approach was adapted from previously reported computational drift-correction and overlay procedures for consecutive AFM force maps^38^.

### Bell-Evans dynamic force spectroscopy

Unbinding force as a function of loading rate was fitted to

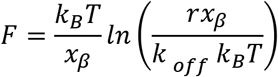

where r is the loading rate, xβ is the distance to the transition state, and koff is the zero-force dissociation rate; koff and xβ were obtained from the intercept and slope of F versus ln(r). Activation energy was estimated from

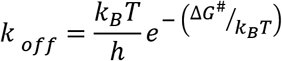

### Worm-like chain fitting and stretching modulus

Force-extension curves with specific adhesion were fitted with the WLC model,

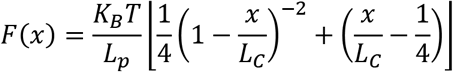

with persistence length (Lp, varied 1–8 nm) and contour length (Lc) as fit parameters, using FODIS software^20^. Only curves showing polymer-like stretching with R² > 0.96 were retained; the 100 best-fitted curves per group were used to calculate stretching modulus,

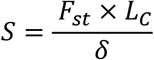

where Fst is the stretching force, and δ is the extension at rupture.

### TGF-β1-induced EMT

MCF-7 cells were treated with recombinant TGF-β1 (10 ng mL⁻¹; PeproTech, Cat#100-21), and metaphase chromosomes were isolated and analyzed by AFS as described above.

### Centromeric proximity mapping and cell line-specific DNA methylation analysis of EMT-associated genes

Centromeric coordinates for the GRCh38 human reference genome assembly were retrieved from the UCSC Genome Browser database and merged per chromosome into unified centromeric regions (minimum Start to maximum End coordinates). Protein-coding gene annotations were obtained from NCBI RefSeq for GRCh38 human genome. For each gene in the Hallmark epithelial-mesenchymal transition gene set, the signed positional distance (upstream/downstream, Mb) was calculated from its gene coordinates (Start and End) relative to the merged centromeric region of its resident chromosome^23^ (**Fig. S1A**) DNA methylation levels of the same gene set in MDA-MB-231, MCF-7, MDA-MB-468, and MCF-10A were obtained from a publicly available methylation array dataset (GEO accession GSE42944), averaged per gene per cell line, and expressed as a Z-score across the panel (**Fig. S1B**). A directly comparable public H3K9me3 dataset with adequate centromeric/pericentromeric coverage across all four lines was not identified; H3K9me3 was therefore assessed only by the AFM-based measurements described above.

### Machine-learning classification

Four supervised classifiers - distance-weighted k-nearest neighbors (weighted k-NN), support vector machine (SVM), random forest, and Gaussian naive Bayes - were evaluated in MATLAB R2025a across five experiments (**Table S6**). Experiment E1 classified pooled cancer observations (MCF-7, MDA-MB-231, and MDA-MB-468) against non-tumorigenic MCF-10A using all 11 features: centromere height, width, and roughness; 5mC- and H3K9me3-specific binding probability, contour length, stretching modulus, and unbinding distance. Experiments E2 and E3 performed three-class differentiation of MCF-7, MDA-MB-231, and MDA-MB-468 using, respectively, the three morphological descriptors and the eight non-colocalized molecular-mechanical descriptors. Experiments E4 and E5 differentiated MDA-MB- 231 from MDA-MB-468 using, respectively, the eight non-colocalized features and a seven-feature colocalized representation comprising colocalized binding probability together with marker-specific contour length, stretching modulus, and unbinding distance. Each cell line contributed 30 chromosome-level observations. Performance was estimated using 10 independently seeded repetitions of stratified nested cross-validation. Each repetition used five outer folds for performance estimation and four inner folds within every outer-training partition for hyperparameter selection by macro-F1. Identical outer partitions were used across classifiers within each experiment. For weighted k-NN and SVM, z-score parameters were estimated exclusively from the relevant training partition and then applied to the validation or test partition. Uniform class priors were used. The five outer-fold predictions were concatenated within each repetition before calculating accuracy, balanced accuracy, class-specific precision, recall and F1, macro-F1, macro-AUC, and multiclass Matthew’s correlation coefficient. Results are reported as mean ± SD across the 10 repetitions; because the repetitions reuse the same observations, the SD represents resampling variability rather than independent biological replication.

### Statistics and reproducibility

Group comparisons used unpaired t-tests, and all force and mechanical parameters are reported as mean ± SD across the ∼10 number of independent chromosomes from 3 independent biological preparations.

## Data availability

The data will be available on request.

## Code availability

The Python 3 drift-correction based on affine transformation and map-registration routine and the MATLAB R2025a machine-learning classification pipeline used in this study will be deposited in a public code repository with a Zenodo-archived release and DOI.

## Acknowledgements

T.R. thanks SNF season 3 collaboration grant (Collab 0011), Anusandhan National Research Foundation (ANRF/ARG/2025/005072/LS), and the Shiv Nadar Institution of Eminence, for funding and the MFP-3D AFM facility. We thank Dr Jayanta Bhattacharya (IIT Delhi) for providing the MDA-MB-468 cell line. We are grateful to Dr Sourav Mishra (POSTECH, South Korea) for the discussion on AFM image drift correction. M.K.J. is supported by Param Hansa Philanthropies.

## Author contributions

TA: investigation, data analysis, first draft. SC: machine-learning analysis. SK, SS and MKJ: computational data analysis. DP and SAK: data analysis. SM and SM: Python-based drift correction. T.R. design, funds acquisition, supervision, and draft editing.

---

**Figure S1:**
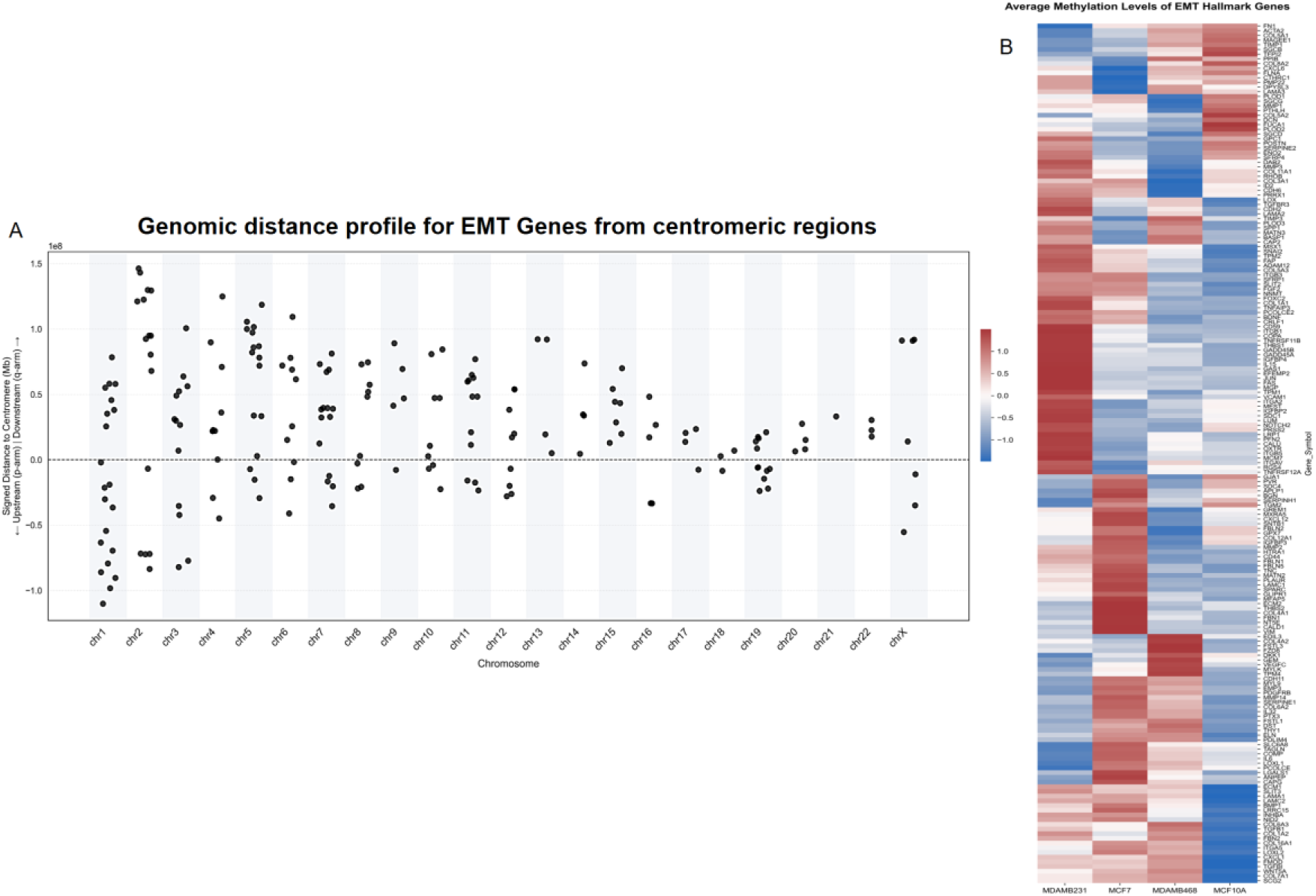
Genomic distribution and DNA methylation patterns of EMT genes across breast cell lines. (A) Genomic distance of EMT-associated genes from their respective centromeric regions across human chromosomes. Distances are shown as signed values, representing the relative position of each gene upstream or downstream of the corresponding centromere. (B) Heatmap showing the average DNA methylation levels of EMT-associated genes in MDA-MB-231, MCF-7, MDA-MB-468, and MCF10A cells based on publicly available methylation data (GSE42944). Methylation values are represented as Z-scores, with positive and negative values indicating relatively higher and lower methylation levels, respectively. **NOTE:** A limitation of the present analysis is the limited availability of reasonably complete, directly comparable publicly available H3K9me3 data in all four breast cell lines analyzed. While publicly available DNA methylation and centromere-gene distance data are available, they were not sufficiently consistent in H3K9me3 signal coverage/quality for a robust comparative analysis of the datasets across MCF10A, MCF-7, MDA-MB-231, and MDA-MB-468 chromosomes, particularly regarding centromeric/pericentromeric activity. Consequently, a quantitative comparison of publicly available data was omitted, and the interpretations are limited to the experimentally measured H3K9me3 levels obtained from AFM-based analysis. High-depth and centromere-resolved H3K9me3 data across the four cell lines are required to further solidify the chromosomal genomic context of the epigenetic alterations reported here.

**Figure S2.**
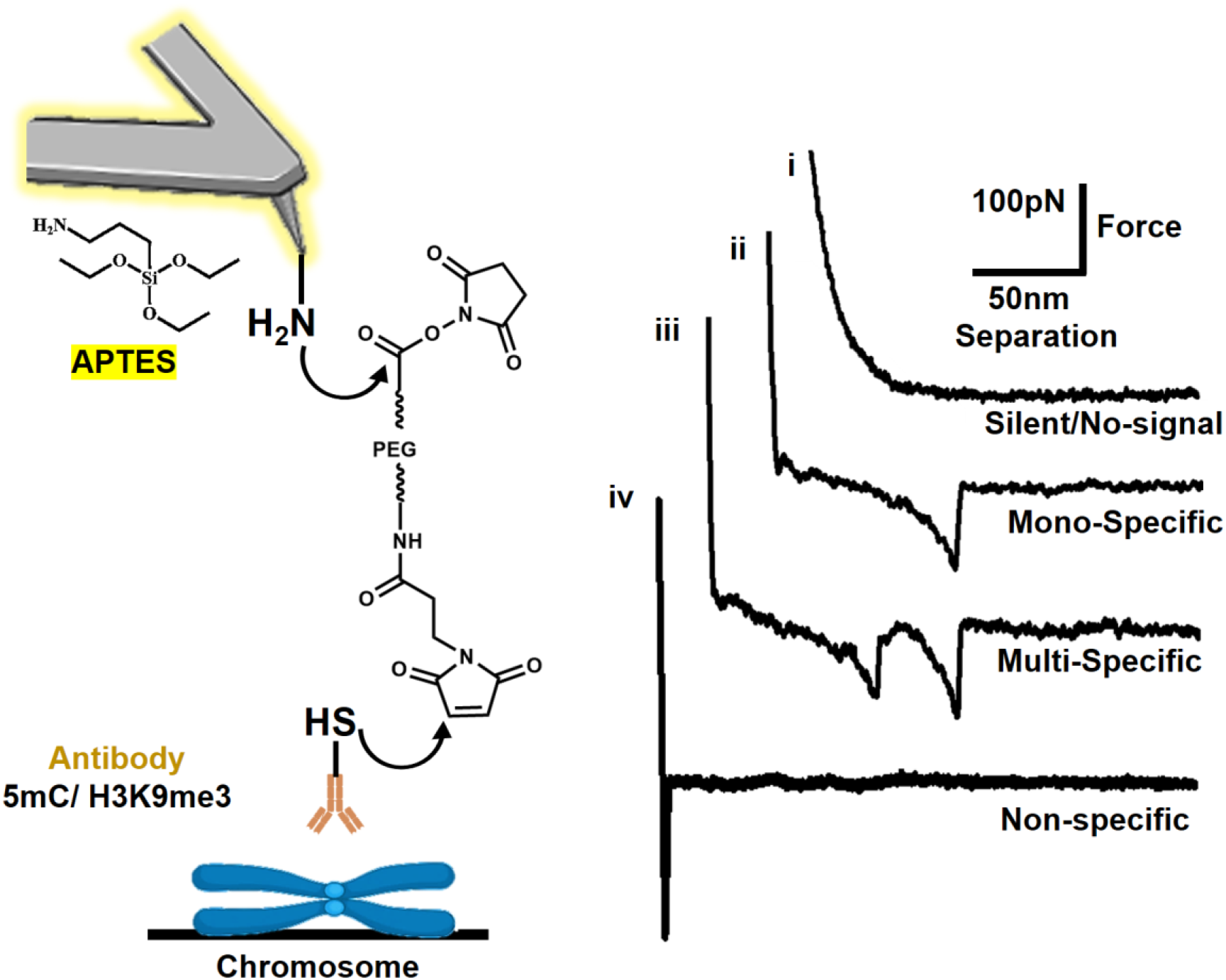
Schematic representation of AFM probe functionalization and representative force-distance curves illustrating specific and nonspecific unbinding events. During the AFM unbinding experiment, the functionalized probe may not encounter a cognate epitope on the chromosome surface, resulting in a force-distance curve without a detectable adhesion event (curve i, inset). Upon specific recognition and binding of the antibody-functionalized probe to its corresponding epigenetic target, a characteristic nonlinear rupture event is observed during the retraction segment of the curve (curves ii and iii), referred to as a specific unbinding event. In contrast, nonspecific interactions produce distinct force signatures that can be differentiated from specific antibody-epitope interactions (curve iv), enabling reliable identification of specific recognition events.

**Scheme S1:**
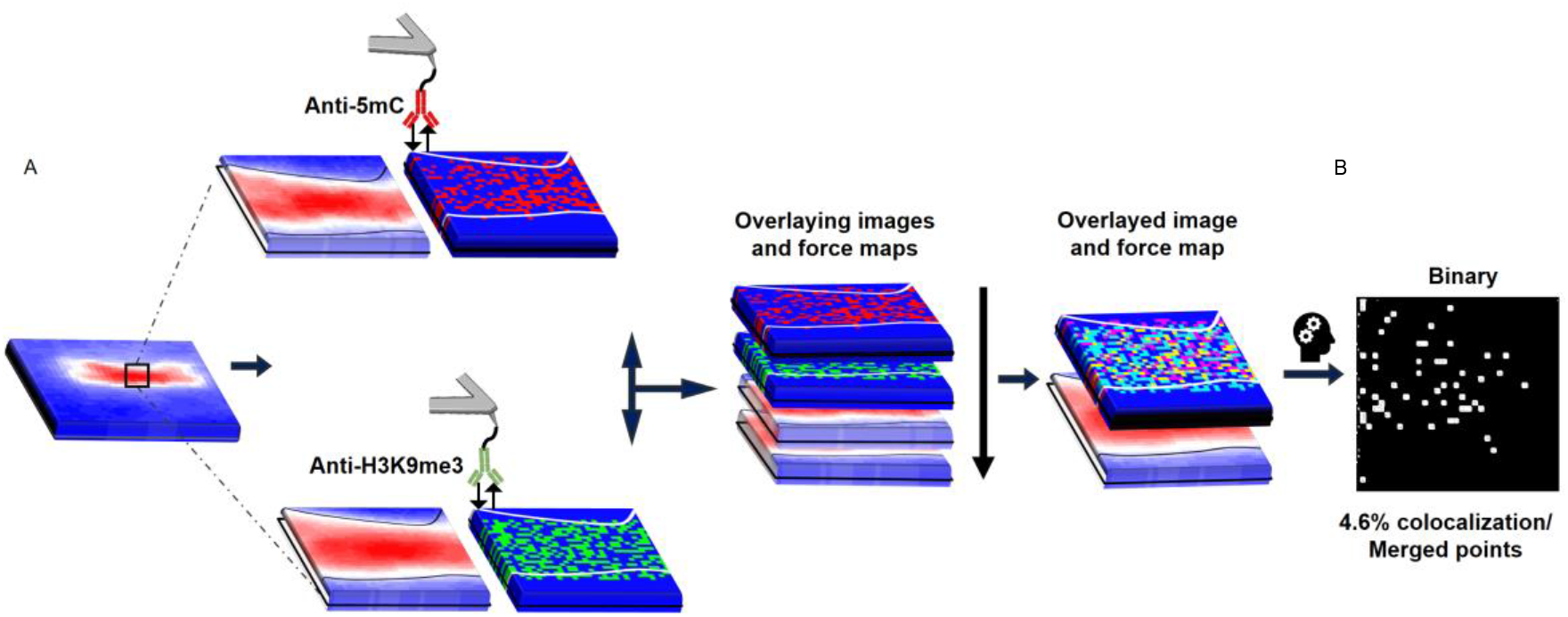
AFM-based dual mapping 5mC and H3K9me3 at chromosome centromeres. (A) Schematic representation of nanoscale mapping of a centromere using AFM tips functionalized with specific antibodies, anti-5mC and anti-H3K9me3 probes, which selectively recognize methylated DNA (red) and histone H3 lysine 9 trimethylation (green), respectively. The resulting force maps are subsequently overlaid to generate an overlayed force map, enabling visualization of the spatial distribution and nanoscale co-localization of 5mC and H3K9me3. (B) ImageJ-based quantification of 5mC/H3K9me3 co-localization. The overlaid force map was imported into ImageJ and converted into a binary image using the Make Binary function. The co-localized points (yellow) were counted, and their percentage was calculated relative to the total number of pixels in the mapped region. Map size: 500 × 500 nm² (32 × 32 pixels).

**Figure S3.**
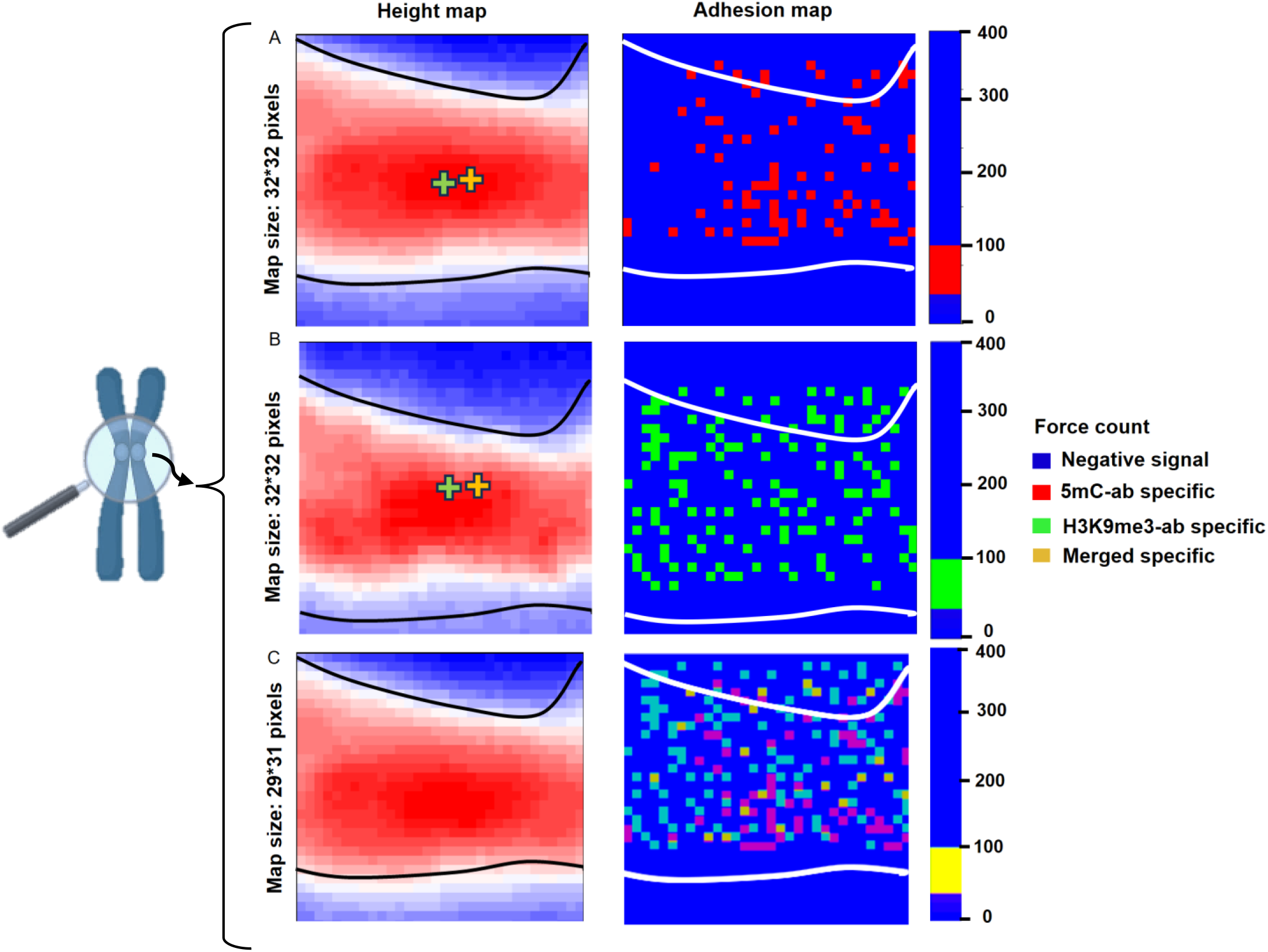
Force-based dual mapping of a centromeric region of a chromosome with AFM tips tethered to (A) Anti-5mC and (B) Anti-H3K9me3 specific antibodies. (C) Drift-corrected overlaid maps of a centromeric region highlighting the co-localization and dual detection of 5mC and H3K9me3 sites. (green mark: geometric centroid; yellow mark: centromere centroid) Drift correction coordinates (translation values) used for alignment: (-2.39, 26.74) (this centromere is from MDA-MB-468 chromosomes) **NOTE: Drift-Corrected Overlay of AFM Force Maps at the Centromeric Region** To improve the spatial correspondence between consecutively acquired AFM force maps, we collected sequential maps from the same centromeric region of an individual chromosome and corrected for lateral drift before overlay by applying basic affine transformations (translation and rotation); based on the co-ordinates of two reference points. The affine transformations are performed on the whole image using custom Python 3 code. In the AFM force-mapping mode, the corresponding height and force information were recorded simultaneously and converted into two-dimensional images. The geometric centroid of the chromosome was extracted from the height distribution of each map and used as a positional reference for registration. Two consecutive maps were treated as two image layers, with the first map kept fixed while the second map was translated pixel-by-pixel along the x and y directions. For each displacement, the mismatch between the geometric centroids (treated as the two reference points for the affine transformation) of the two maps was calculated. The x–y displacement corresponding to the minimum centroid mismatch was considered the lateral drift between the consecutive maps. The calculated displacement (translation and rotation of the two reference points) was then applied to the corresponding force map to align it with the reference map. The same procedure was repeated for subsequent maps to obtain the drift-corrected overlay. Representative height maps and corresponding force maps from the centromeric region are shown before and after registration. The corrected overlays show improved spatial correspondence of the force-map features within the centromeric region, allowing the molecular interaction signals detected in consecutive acquisitions to be compared at the same spatial coordinates.

**Figure S4.**
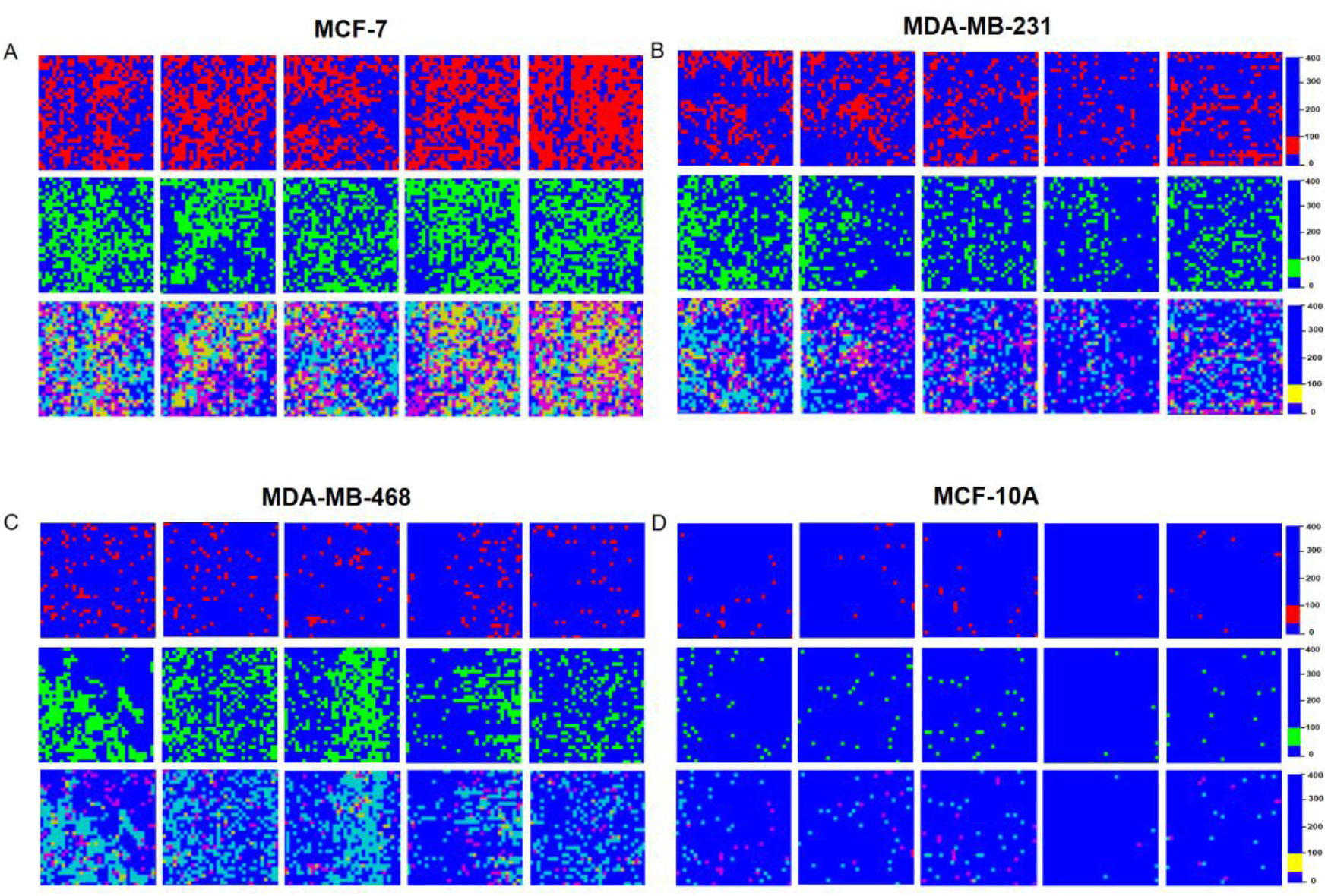
Five representative force maps of DNA, histone methylation and their co-localized marks of (A) MCF-7, (B) MDA-MB-231, (C) MDA-MB-468 and (D) MCF-10A chromosome centromeres, demonstrating the reproducibility and spatial consistency of epigenetic mark distribution across independent measurements.

**Figure S5:**
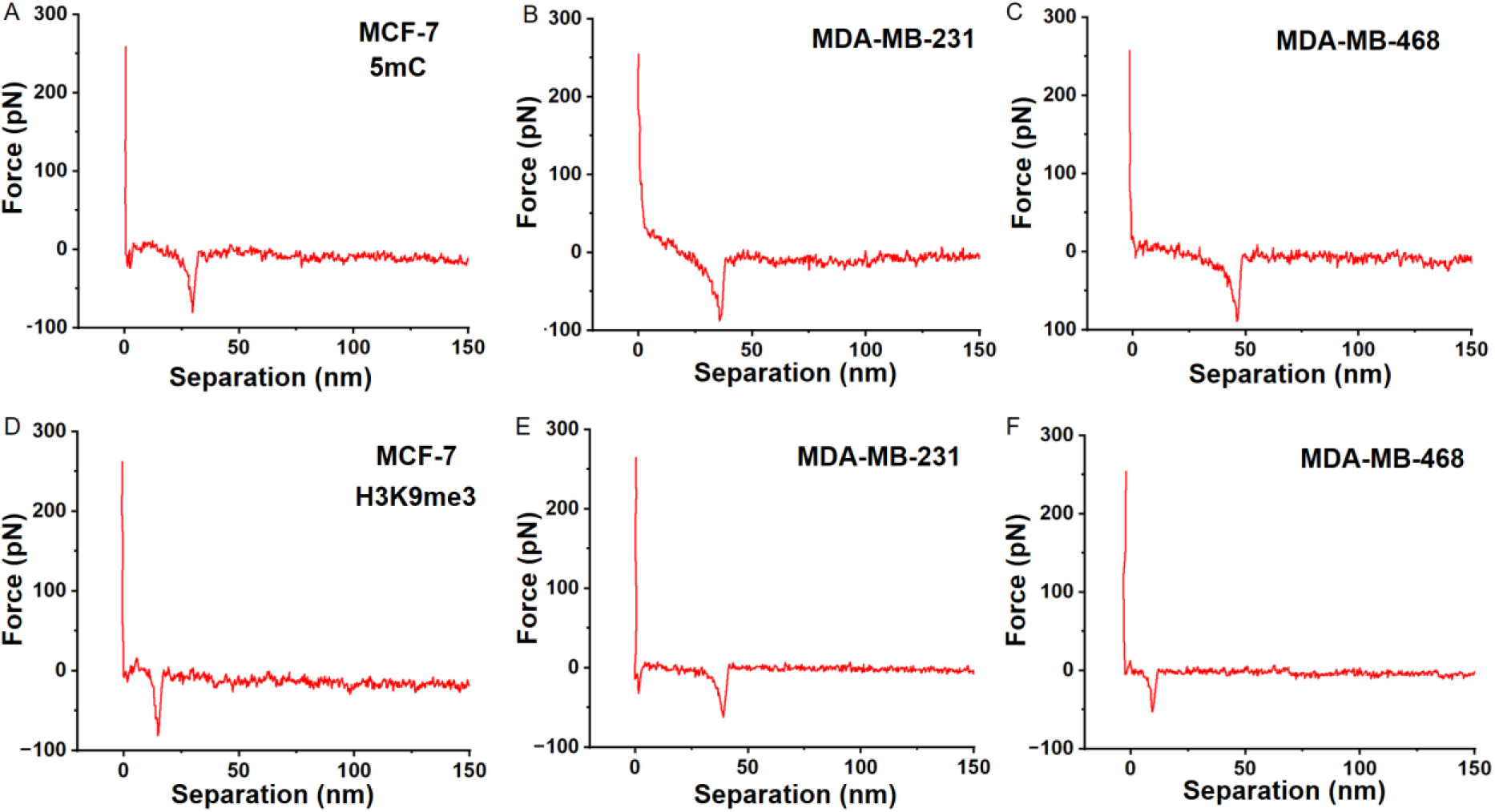
Representative force-separation curves of AFS measurements of each chromosome for both 5mC and H3K9me3 interactions.

**Figure S6.**
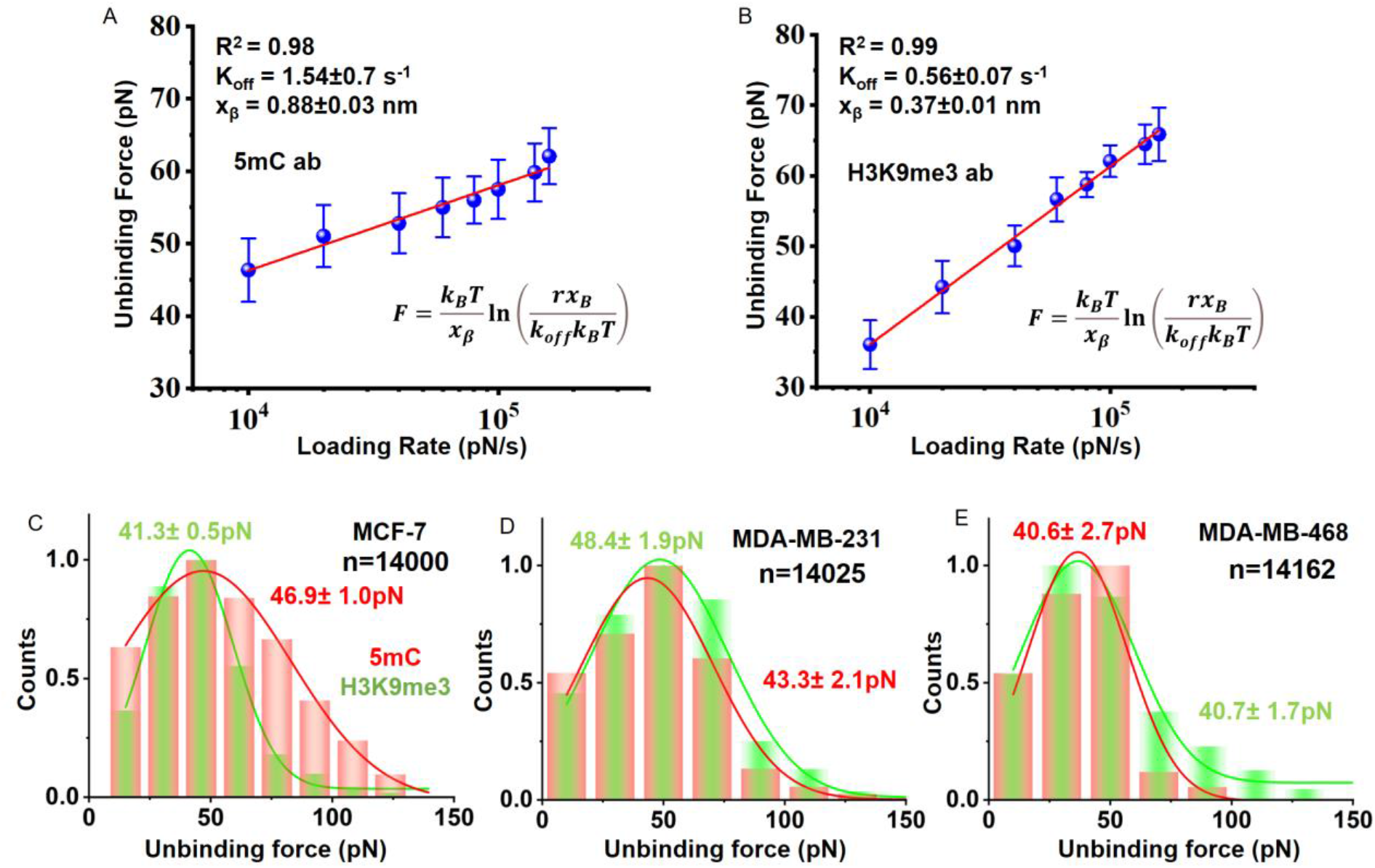
Dependence of the unbinding force between (A) anti-5mC and (B) anti-H3K9me3 antibodies and methylated site on the logarithm of loading rate. The blue spheres indicate data points for both cases, respectively. The solid red lines represent the numerical fit of the experimental data with Bell-Evans model. The obtained kinetic parameters are shown in the inset. Histograms of the unbinding force distributions for (C) MCF-7, (D) MDA-MB-231 and (E) MDA-MB-468 chromosomes for both 5mC and H3K9me3 interactions, respectively. The mean unbinding force values were determined from the single-peak Gaussian distributions.

**Figure S7.**
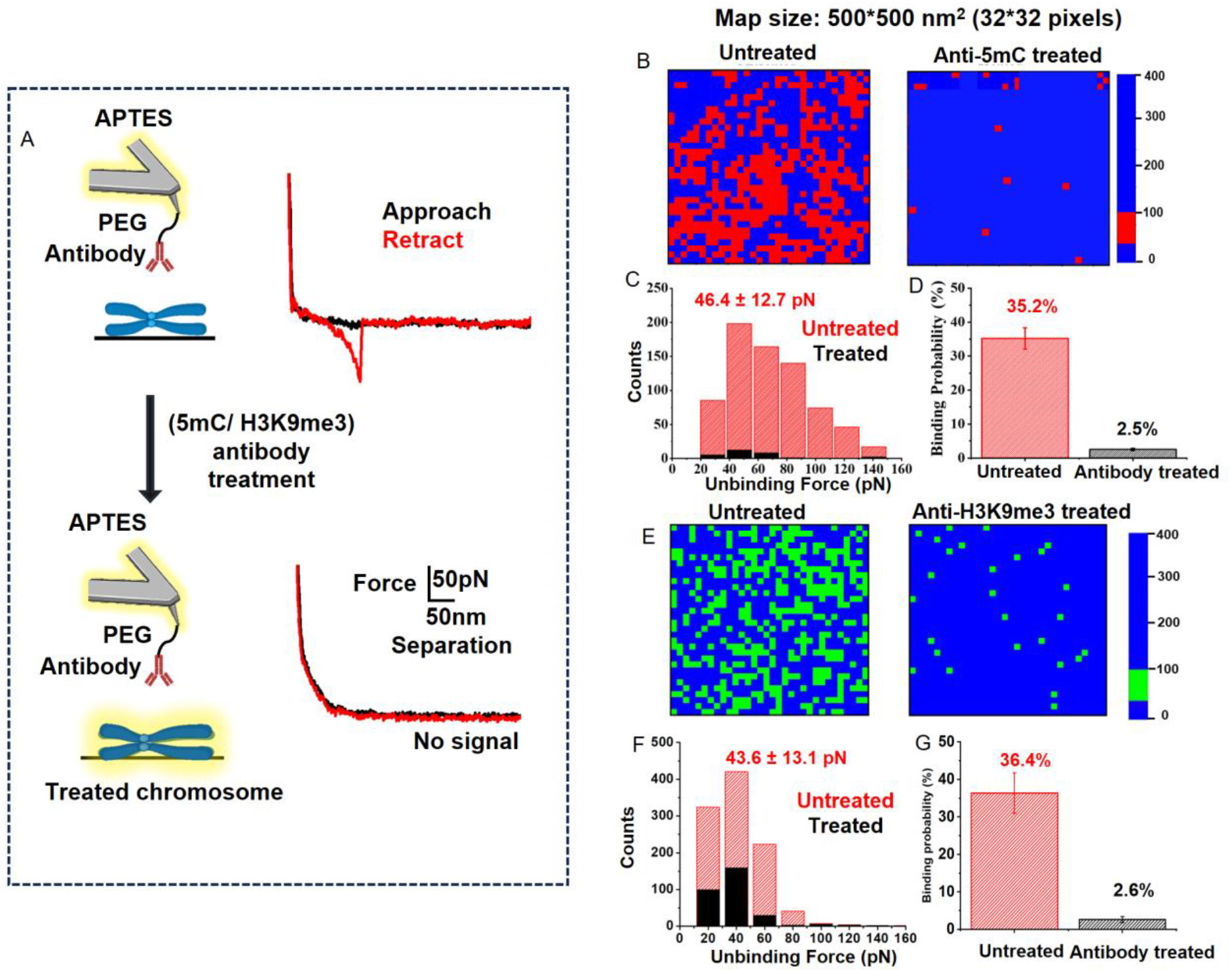
Validation of specific interactions between antibodies and methylated sites. (A) Representative force curve obtained using an AFM tip functionalized with a specific antibody probing a methylated chromosome. The approach and retraction curves are shown in black and red, respectively. The absence of adhesion events after blocking the methylated sites with free antibodies confirms interaction specificity. Force maps showing the spatial distribution of 5mC (B) and H3K9me3 (E) before and after blocking. Histograms of unbinding force distributions for 5mC (C) and H3K9me3 (F) interactions before (red) and after (black) blocking, demonstrating a significant reduction in specific binding events. Quantitative analysis of binding probabilities for 5mC (D) and H3K9me3 (G) before and after blocking, further confirming the specificity of antibody-methylation interactions.

**Figure S8.**
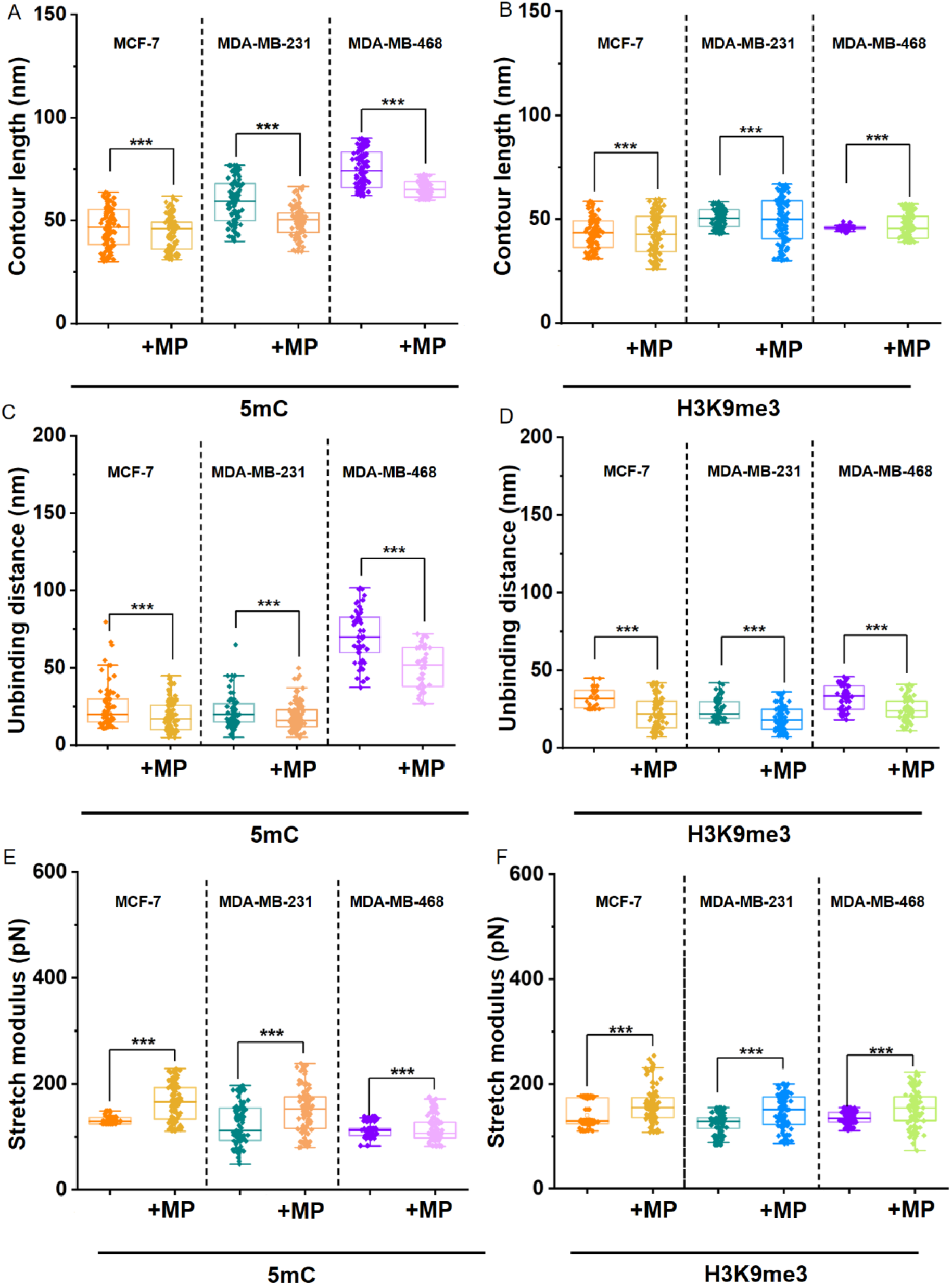
Box plot analysis of mechanical parameters for single and co-localized interactions. (A, B) Contour length, (C, D) unbinding distance, and (E, F) stretch modulus for 5mC and H3K9me3 interactions across all chromosomes. The plots highlight the statistical differences between single-binding events and co-localized interaction points (MP: Merged points/co-localized sites).

**Figure S9.**
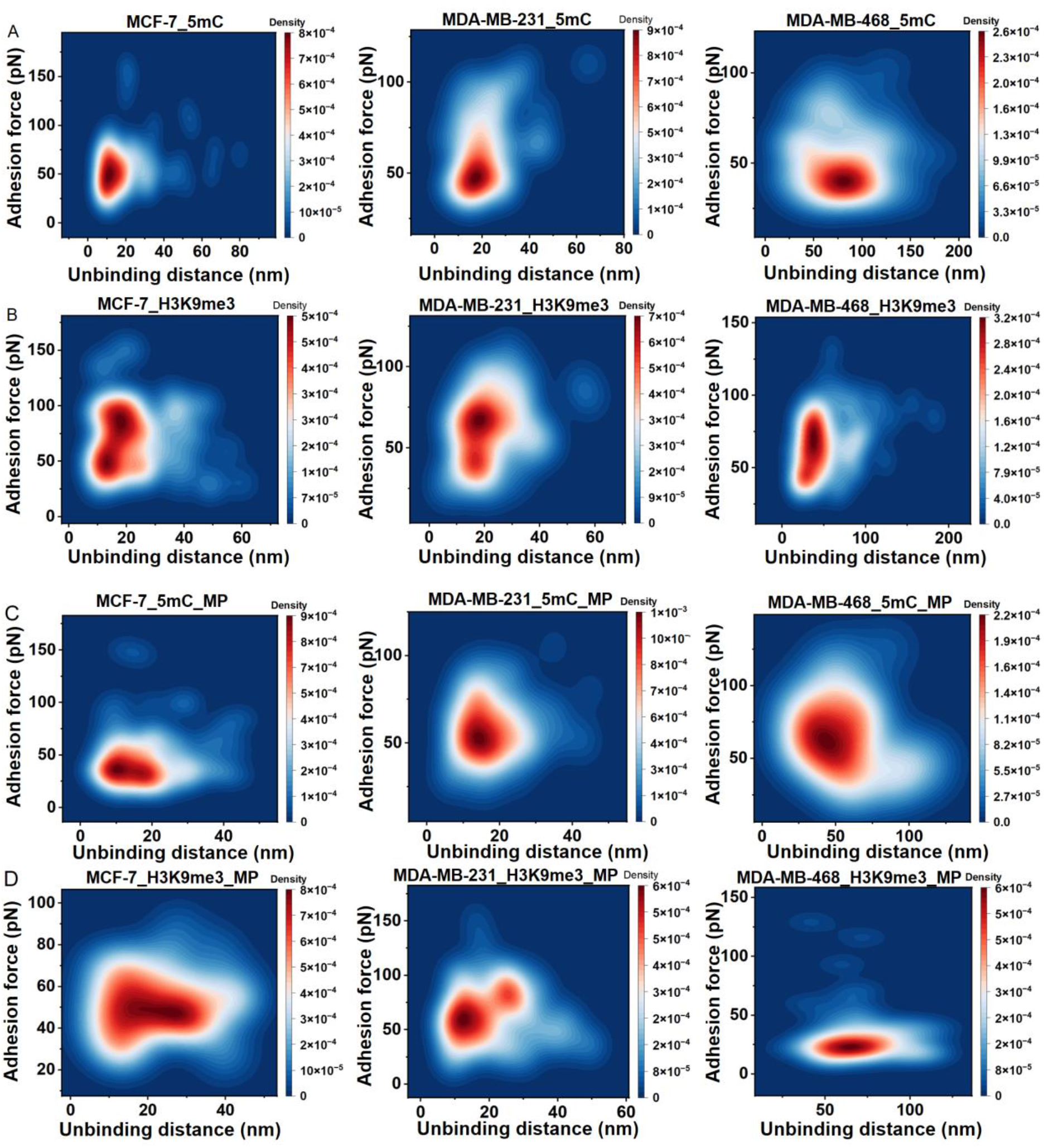
2D Kernel density estimation (KDE) contour maps of force-distance interactions between antibodies and methylated chromosomes. KDE contour maps representing the distribution of unbinding events for 5mC (A) and H3K9me3 (B) antibodies interacting with MCF-7, MDA-MB-231, and MDA-MB-468chromosomes. Corresponding KDE maps for co-localized (MP: Merged points) interaction points for 5mC (C) and H3K9me3 (D) across the same cell lines. Each contour map represents the density of unbinding events across unbinding distance and force, as obtained from AFM-based measurements. Denser regions (brighter contours) indicate frequent binding events, enabling direct comparison of interaction profiles and heterogeneity across different chromosome types and interaction modes.

**Figure S10.**
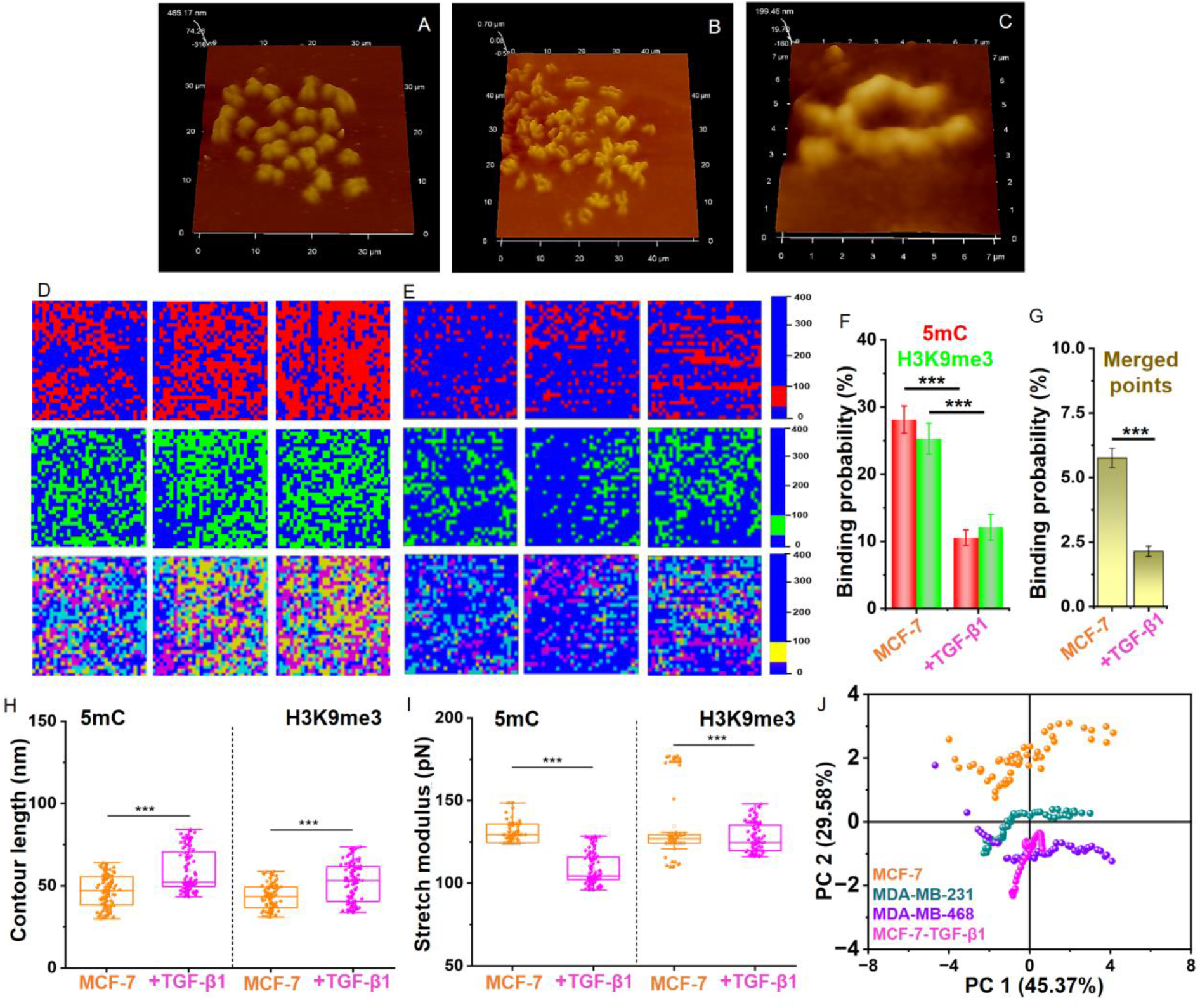
AFM-based recognition mapping reveals TGF-β1-induced epigenetic remodeling of metaphase chromosomes. Representative AFM topography images of metaphase chromosomes isolated from (A) untreated MCF-7 cells and (B, C) TGF-β1-treated MCF-7 cells. High-resolution Force maps (500*500nm^2^;32*32 pixels) acquired from the centromeric regions of chromosomes isolated from (D) MCF-7 and (E) TGF-β1-treated MCF-7 chromosomes. Percentages of Binding probabilities occupied by the positive pixel (F) for 5mC or H3K9me3 and for merged points of both chromosomes. Box plots showing the mechanical parameters for both interactions (H) Contour length and (I) stretch modulus for 5mC and H3K9me3 interactions across both chromosomes. (J) Principal component analysis (PCA) of AFM-derived recognition parameters comparing untreated MCF-7, MDA-MB-231, MDA-MB-468, and TGF-β1-treated MCF-7, chromosomes. TGF-β1-treated MCF-7 chromosomes shift toward the TNBC cluster, indicating EMT-associated epigenetic remodeling at the chromosome scale. Statistical significance was determined using an unpaired Student’s t-test; p < 0.05.

**Figure S11.**
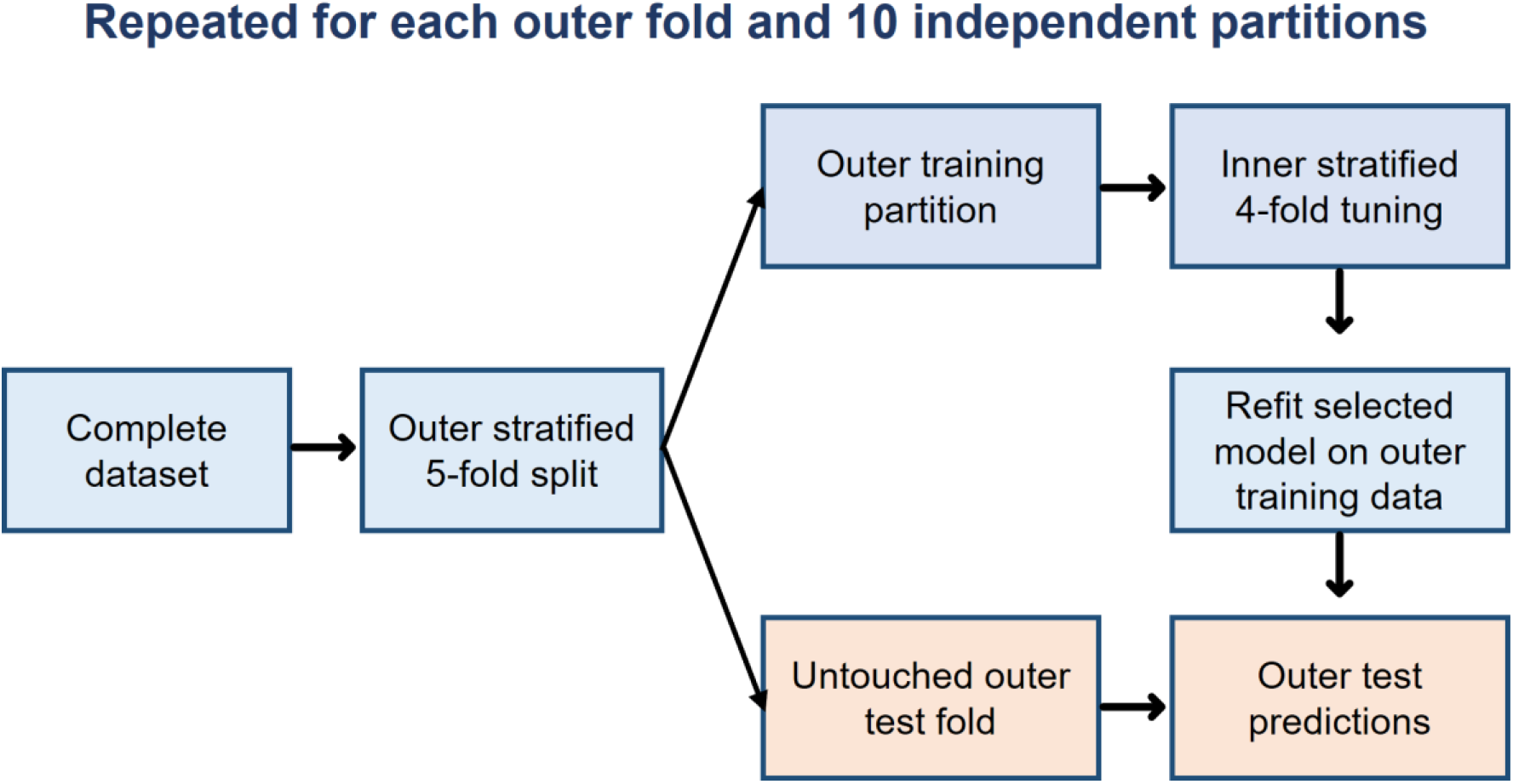
Repeated nested cross-validation workflow. The inner loop selected model hyperparameters, whereas the outer loop estimated generalization performance. All data-dependent preprocessing was fitted using training folds only.

**Figure S12.**
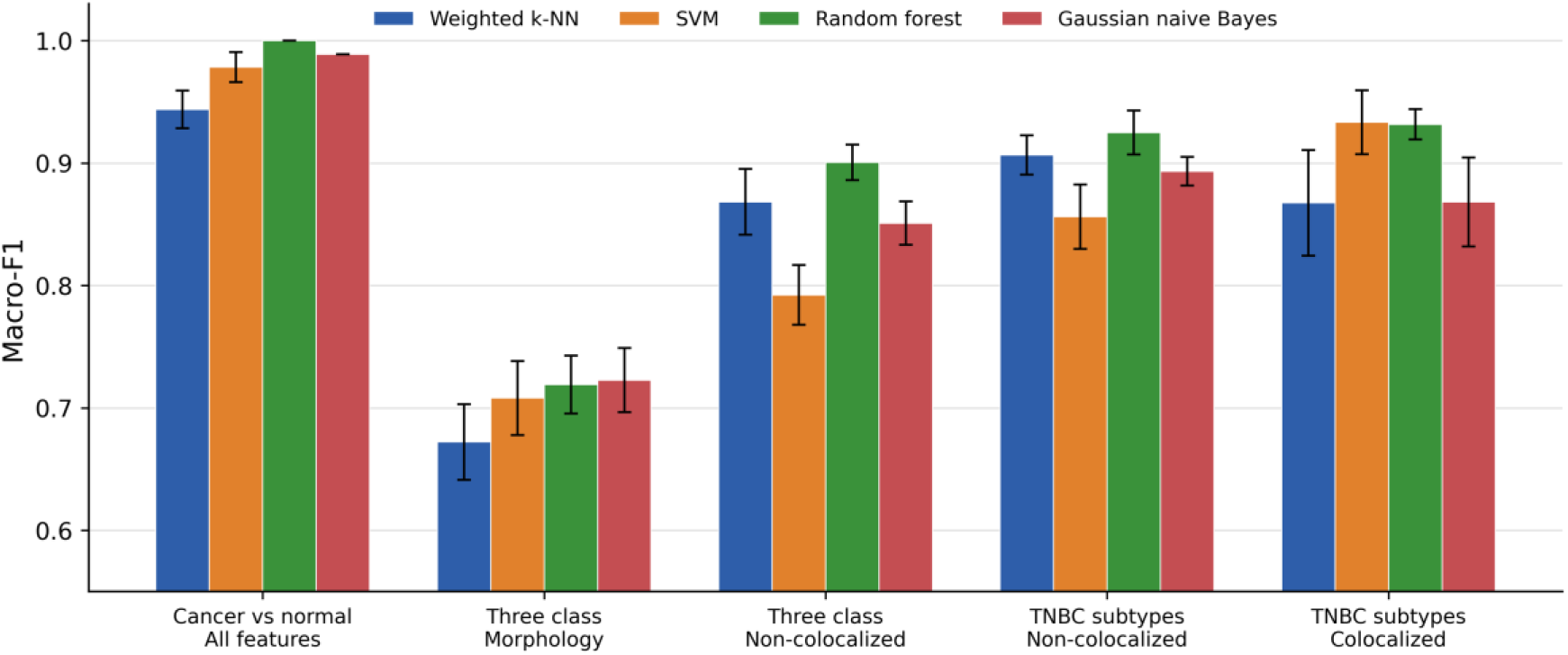
Macro-F1 across the five experiments. Bars show the mean across 10 repeated nested cross-validation partitions and error bars show one standard deviation

**Figure S13.**
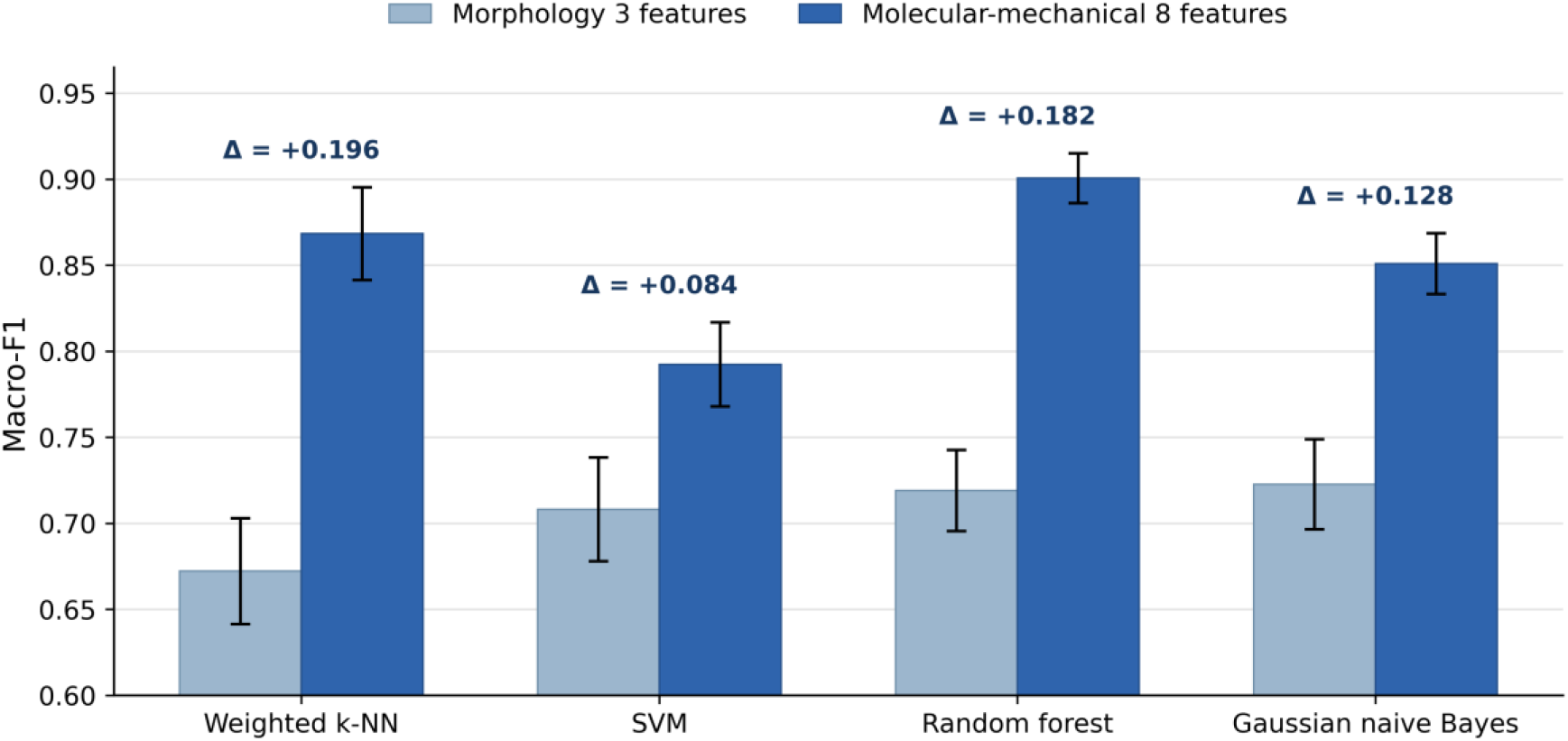
Comparison of morphology and non-colocalized molecular-mechanical features for three-class cancer cell-line differentiation. Numeric annotations show the change in mean macro-F1 after replacing morphology with the eight-feature representation.

**Figure S14.**
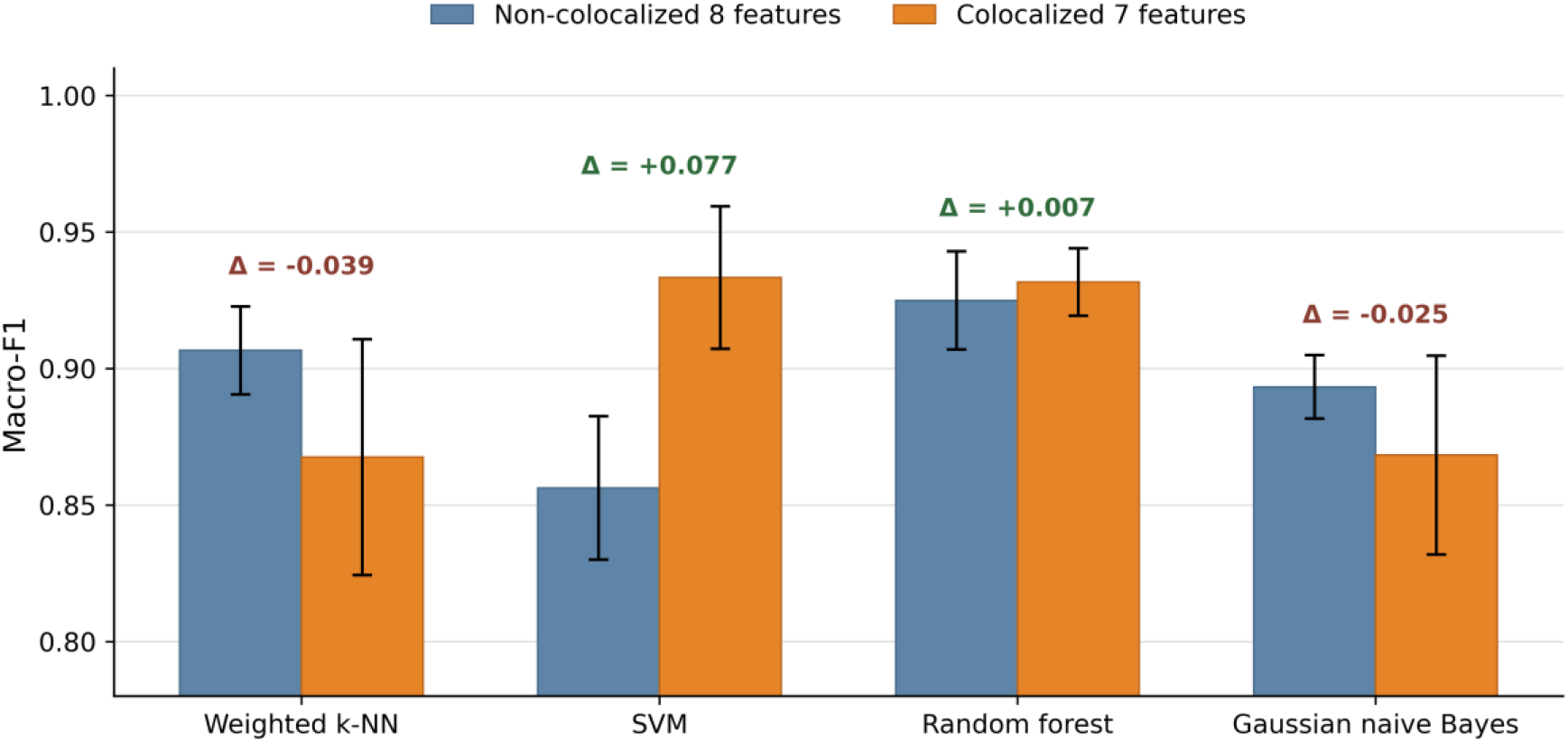
Unpaired descriptive comparison of TNBC subtype classification using the non-colocalized and colocalized datasets. Numeric annotations show the colocalized minus non-colocalized change in mean macro-F1 for each classifier.

**Table S1.** Secondary Structure calculation from CD Spectra of all chromosomes at room temperature.

|  | Secondary structure (%) |  |  |  |  |
| --- | --- | --- | --- | --- | --- |
| Histones | $\alpha$ helix | Antiparallel | Parallel | Turn | others |
| **MCF-7** | 3.6 | 9.6 | 41.8 | 0 | 45 |
| **MDA-MB-231** | 26.2 | 5 | 0 | 8.2 | 65.3 |
| **MDA-MB-468** | 19.4 | 6 | 3.3 | 2.5 | 62.1 |
| **MCF-10A** | 29.7 | 1.6 | 0 | 9.4 | 62.3 |

**Table S2.** FT-IR Peak Assignments.

| Peak Number | Wavenumber (cm<sup>-1</sup>) |  |  |  | Corresponding vibrational modes |
| --- | --- | --- | --- | --- | --- |
|  | MCF-7 | MDA-MB-231 | MDA-MB-468 | MCF-10A |  |
| 1 | 1038 | 1038 | 1038 | 1039 | P=O phospholipid stretching, C-O-C stretching |
| 2 | 1079 | 1076 | 1076 | 1072 | Phosphate symmetric stretching |
| 3 | 1122 | 1120 | 1112 | 1118 | Sugar-phosphate vibrations, Phosphate backbone |
| 4 | 1238 | 1239 | 1240 | 1245 | C=O inorganic carbonate stretching, P=O asymmetric stretching |
| 5 | 1406 | 1406 | 1407 | 1406 | Antisymmetric aliphatic C-H deformation of methyl and methoxyl |
| 6 | 1594 | 1593 | 1590 | 1595 | C-N Amide group stretching, NH bending |
| 7 | 1663 | 1659 | 1659 | 1661 | C=O Amide group (Cytosine) |
| 8 | 1710 | 1710 | 1713 | 1715 | C=O stretching (carboxylic group) |
| 9 | 2884 | 2884 | 2884 | 2885 | C-H or CH<sub>2</sub> Symmetric Stretching vibrations |
| 10 | 2941 | 2941 | 2941 | 2940 | CH<sub>3</sub> vibration or C-H- Antisymmetric Stretching vibrations |
| 11 | 3341 | 3350 | 3348 | 3349 | O-H stretching /H-bonding |

**Table S3.** Statistical Information for DNA-CH3 Band Peak and Area Under the Curve Values for DNA Obtained from FT-IR Spectral Profiles.

| DNA | Peak starts (cm<sup>-1</sup>) | Peak ends (cm<sup>-1</sup>) | Area units | Ratio cancer/healthy |
| --- | --- | --- | --- | --- |
| MCF-7 | 2850 | 2987 | 0.88 | 1.5 |
| MDA-MB-231 | 2858 | 2981 | 0.73 | 1.2 |
| MDA-MB-468 | 2860 | 2984 | 0.64 | 1.1 |
| MCF-10A | 2863 | 2981 | 0.58 | - |

**Table S4.** Summary of average structural, mechanical, and interaction parameters of chromosomes, including height, width, surface roughness, unbinding force, binding probability, contour length, and stretch modulus across all studied cell lines.

| Parameter<br>(n= no. of chromosomes) | Non-TNBC<br>(MCF-7) |  | TNBC; Basal B<br>(MDA-MB-231) |  | TNBC; Basal A<br>(MDA-MB-468) |  | Control<br>(MCF-10A) |  |
| --- | --- | --- | --- | --- | --- | --- | --- | --- |
|  | 5mC | H3K9me3 | 5mC | H3K9me3 | 5mC | H3K9me3 | 5mC | H3K9me3 |
| Height (nm) (n=50) | 141.1±13.1 |  | 160.9±15.3 |  | 194.3±42.4 |  | 181.8±12.6 |  |
| Width (µm) (n=50) | 1.8±0.5 |  | 1.5±0.4 |  | 1.4±0.3 |  | 2.0±0.6 |  |
| Roughness (nm) (n=50) | 72.6±12.3 |  | 94.7±17.4 |  | 114.36±29.4 |  | 163.7±15.2 |  |
| Unbinding Force (pN)<br>(n=10) | 46.9±1.0 | 41.3±0.5 | 43.3±2.1 | 48.4±1.9 | 40.6± 2.7 | 40.7± 1.7 | - | - |
| Binding Probability (%)<br>(n=10) | 28.2±2.1 | 25.2±2.2 | 18.1±4.2 | 12.9±2.0 | 11.7±1.1 | 18.7±1.2 | 2.1±0.6 | 3.7±0.4 |
| Contour length (nm)<br>(n=10) | 41.7±8 | 36.2±7 | 48.7±11 | 47.5±12 | 78.3±21 | 43.4±14 | - | - |
| Stretch Modulus (pN)<br>(n=10) | 124.6±6 | 130.7±17 | 90.1±32 | 110.3±5 | 102±7 | 128.7±18 | - | - |
| Colocalized points<br>Binding Probability (%)<br>(n=10) | 5.7±0.3 |  | 2.1±0.6 |  | 1.5±0.3 |  | 0.4±0.08 |  |
| Colocalized points<br>Contour length (nm)<br>(n=10) | 32.1±9 | 39.9±10 | 44.8±14 | 44.7±9 | 67.7±13 | 45.8±15 | - | - |
| Colocalized points<br>Stretch Modulus (pN)<br>(n=10) | 166.4±33 | 159.3±32 | 149.2±36 | 149±30 | 94.5±10 | 150.4±33 | - | - |

**Table S5.** Summary of dissociation kinetic parameters for interactions between anti-5mC and anti-H3K9me3 antibodies with chromosomes.

| Parameter | anti-5mC | anti-H3K9me3 |
| --- | --- | --- |
| Dissociation rate constant<br>(s<sup>-1</sup>) | 1.54±0.7 | 0.56±0.07 |
| Dissociation energy barrier<br>(nm) | 0.88±0.03 | 0.37±0.01 |
| Activation energy (kcal/mol) | 17.1 | 17.78 |
| Life-time (s) | 0.65 | 1.78 |

**Table S6.** Classification tasks, sample sizes and feature representations of Machine learning analysis.

| ID | Task | Classes | N | Feature set |
| --- | --- | --- | --- | --- |
| E1 | Cancer vs normal | Cancer: MCF-7, MB-231, MB-468; normal: MCF-10A | 120 | All 11 features |
| E2 | Three-class cancer differentiation | MCF-7 vs MB-231 vs MB-468 | 90 | Height, width, roughness |
| E3 | Three-class cancer differentiation | MCF-7 vs MB-231 vs MB-468 | 90 | 8 non-colocalized features |
| E4 | TNBC subtype differentiation | MB-231 vs MB-468 | 60 | 8 non-colocalized features |
| E5 | TNBC subtype differentiation | MB-231 vs MB-468 | 60 | 7 colocalized features |

**Table S7.**
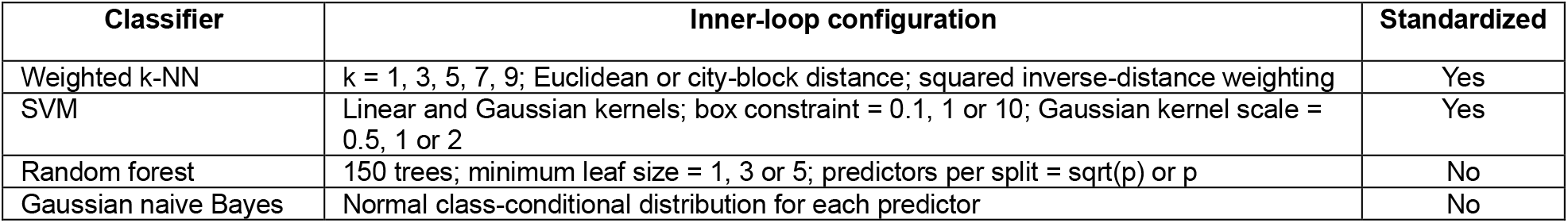
Classifier configurations evaluated within the inner cross-validation loop.

**Table S8.** Repeated nested cross-validation performance for Experiments E1 to E3.

| Exp. | Classifier | Accuracy | Balanced acc. | Macro-F1 | Macro-AUC | MCC |
| --- | --- | --- | --- | --- | --- | --- |
| E1 | Weighted k-NN | 0.956 ± 0.012 | 0.964 ± 0.011 | 0.944 ± 0.015 | 0.964 ± 0.011 | 0.891 ± 0.029 |
| E1 | SVM | 0.983 ± 0.010 | 0.989 ± 0.006 | 0.978 ± 0.012 | 0.999 ± 0.001 | 0.958 ± 0.023 |
| E1 | Random forest | 1.000 ± 0.000 | 1.000 ± 0.000 | 1.000 ± 0.000 | 1.000 ± 0.000 | 1.000 ± 0.000 |
| E1 | Gaussian naive Bayes | 0.992 ± 0.000 | 0.983 ± 0.000 | 0.989 ± 0.000 | 1.000 ± 0.000 | 0.978 ± 0.000 |
| E2 | Weighted k-NN | 0.680 ± 0.030 | 0.680 ± 0.030 | 0.672 ± 0.031 | 0.807 ± 0.030 | 0.527 ± 0.045 |
| E2 | SVM | 0.704 ± 0.032 | 0.704 ± 0.032 | 0.708 ± 0.030 | 0.850 ± 0.027 | 0.560 ± 0.050 |
| E2 | Random forest | 0.719 ± 0.025 | 0.719 ± 0.025 | 0.719 ± 0.024 | 0.864 ± 0.015 | 0.580 ± 0.037 |
| E2 | Gaussian naive Bayes | 0.719 ± 0.029 | 0.719 ± 0.029 | 0.723 ± 0.026 | 0.876 ± 0.014 | 0.580 ± 0.042 |
| E3 | Weighted k-NN | 0.869 ± 0.027 | 0.869 ± 0.027 | 0.868 ± 0.027 | 0.955 ± 0.025 | 0.805 ± 0.040 |
| E3 | SVM | 0.792 ± 0.025 | 0.792 ± 0.025 | 0.792 ± 0.024 | 0.930 ± 0.010 | 0.689 ± 0.037 |
| E3 | Random forest | 0.901 ± 0.014 | 0.901 ± 0.014 | 0.901 ± 0.014 | 0.967 ± 0.004 | 0.852 ± 0.021 |
| E3 | Gaussian naive Bayes | 0.851 ± 0.018 | 0.851 ± 0.018 | 0.851 ± 0.018 | 0.953 ± 0.004 | 0.777 ± 0.026 |

**Table S9.** Repeated nested cross-validation performance for Experiments E4 and E5.

| Exp. | Classifier | Accuracy | Balanced acc. | Macro-F1 | Macro-AUC | MCC |
| --- | --- | --- | --- | --- | --- | --- |
| E4 | Weighted k-NN | 0.907 ± 0.016 | 0.907 ± 0.016 | 0.907 ± 0.016 | 0.943 ± 0.023 | 0.814 ± 0.032 |
| E4 | SVM | 0.857 ± 0.026 | 0.857 ± 0.026 | 0.856 ± 0.026 | 0.901 ± 0.028 | 0.718 ± 0.054 |
| E4 | Random forest | 0.925 ± 0.018 | 0.925 ± 0.018 | 0.925 ± 0.018 | 0.975 ± 0.004 | 0.852 ± 0.037 |
| E4 | Gaussian naive Bayes | 0.893 ± 0.012 | 0.893 ± 0.012 | 0.893 ± 0.012 | 0.949 ± 0.005 | 0.788 ± 0.023 |
| E5 | Weighted k-NN | 0.868 ± 0.043 | 0.868 ± 0.043 | 0.868 ± 0.043 | 0.927 ± 0.065 | 0.745 ± 0.083 |
| E5 | SVM | 0.933 ± 0.026 | 0.933 ± 0.026 | 0.933 ± 0.026 | 0.981 ± 0.011 | 0.868 ± 0.052 |
| E5 | Random forest | 0.932 ± 0.012 | 0.932 ± 0.012 | 0.932 ± 0.012 | 0.965 ± 0.005 | 0.864 ± 0.025 |
| E5 | Gaussian naive Bayes | 0.868 ± 0.036 | 0.868 ± 0.036 | 0.868 ± 0.036 | 0.933 ± 0.033 | 0.737 ± 0.073 |

**Table S10.** Class-specific performance of the best-performing model for each experiment.

| Exp. | Selected model | Class | Precision | Recall | F1 | AUC |
| --- | --- | --- | --- | --- | --- | --- |
| E1 | Random forest | Cancer | 1.000 ± 0.000 | 1.000 ± 0.000 | 1.000 ± 0.000 | 1.000 ± 0.000 |
| E1 | Random forest | Normal | 1.000 ± 0.000 | 1.000 ± 0.000 | 1.000 ± 0.000 | 1.000 ± 0.000 |
| E2 | Gaussian naive Bayes | MCF-7 | 0.629 ± 0.039 | 0.627 ± 0.049 | 0.627 ± 0.039 | 0.847 ± 0.014 |
| E2 | Gaussian naive Bayes | MDA-MB-231 | 0.579 ± 0.046 | 0.617 ± 0.042 | 0.596 ± 0.034 | 0.788 ± 0.027 |
| E2 | Gaussian naive Bayes | MDA-MB-468 | 0.979 ± 0.028 | 0.913 ± 0.045 | 0.944 ± 0.028 | 0.994 ± 0.008 |
| E3 | Random forest | MCF-7 | 0.905 ± 0.022 | 0.947 ± 0.036 | 0.925 ± 0.020 | 0.978 ± 0.003 |
| E3 | Random forest | MDA-MB-231 | 0.886 ± 0.026 | 0.847 ± 0.039 | 0.865 ± 0.025 | 0.943 ± 0.008 |
| E3 | Random forest | MDA-MB-468 | 0.914 ± 0.026 | 0.910 ± 0.016 | 0.912 ± 0.015 | 0.981 ± 0.004 |
| E4 | Random forest | MDA-MB-231 | 0.913 ± 0.024 | 0.940 ± 0.038 | 0.926 ± 0.018 | 0.975 ± 0.004 |
| E4 | Random forest | MDA-MB-468 | 0.940 ± 0.036 | 0.910 ± 0.027 | 0.924 ± 0.018 | 0.975 ± 0.004 |
| E5 | SVM | MDA-MB-231 | 0.920 ± 0.026 | 0.950 ± 0.032 | 0.934 ± 0.026 | 0.981 ± 0.011 |
| E5 | SVM | MDA-MB-468 | 0.949 ± 0.033 | 0.917 ± 0.028 | 0.932 ± 0.026 | 0.981 ± 0.011 |

**Table S11.** Descriptive comparison of non-colocalized and colocalized TNBC feature representations.

| Classifier | Non-coloc. F1 | Coloc. F1 | F1 change | Non-coloc. AUC | Coloc. AUC |
| --- | --- | --- | --- | --- | --- |
| Weighted k-NN | 0.907 ± 0.016 | 0.868 ± 0.043 | -0.039 | 0.943 | 0.927 |
| SVM | 0.856 ± 0.026 | 0.933 ± 0.026 | +0.077 | 0.901 | 0.981 |
| Random forest | 0.925 ± 0.018 | 0.932 ± 0.012 | +0.007 | 0.975 | 0.965 |
| Gaussian naive Bayes | 0.893 ± 0.012 | 0.868 ± 0.036 | -0.025 | 0.949 | 0.933 |

